# An Adipose mTORC1 Effector Links Nutrient-induced Ribosomal Protein Translation to Lifespan in *Drosophila*

**DOI:** 10.1101/2025.05.27.656475

**Authors:** Jun Wang, Zixin Cai, Jiaojiao Gu, Meng Yang, Kexin Chang, Xinrui Ning, Yilin Wen, Yan Yan, Jiongming Lu, Yirong Wang, Zongzhao Zhai

**Affiliations:** Hunan Provincial Key Laboratory of Animal Intestinal Function and Regulation, College of Life Sciences, Hunan Normal University, Changsha, China; Hunan Research Center of the Basic Discipline for Cell Signaling, College of Biology, Hunan University, Changsha, China; Division of Life Science, Hong Kong University of Science and Technology, Hong Kong, China; Shanghai Institute of Nutrition and Health, University of Chinese Academy of Sciences, Chinese Academy of Sciences, Shanghai 200031, China; School of Basic Medical Sciences, Hunan Normal University, Changsha, China

## Abstract

Mechanistic target of rapamycin complex 1 (mTORC1) senses nutrient availability to orchestrate metabolic processes critical for physiological homeostasis and organismal ageing^1^. While mTORC1 preferentially regulates the translation of 5′-terminal oligopyrimidine (TOP) motif-containing mRNAs that predominantly encode ribosomal proteins (RPs) via the translational repressor 4E-BP ^2^, this mTORC1 function is resistant to rapamycin inhibition ^3^. TOP mRNAs are exceptionally abundant, thus imposing a major translational burden on cells; yet how their translation is physiologically tuned and linked to longevity remain unexplored. Here we identify Lsp2, originally known as a storage protein ^4^, as an adipose effector of mTORC1 that modulates lifespan in *Drosophila*. Lsp2 is induced by essential amino acids (EAAs) via mTORC1. Genetic ablation of *Lsp2* to blunt organismal response to protein diets drives robust lifespan extension without compromising key life-history traits such as reproduction. Translatomic profiling reveals that loss of *Lsp2* selectively reduces global TOP mRNA translation in a 4E-BP-dependent manner, thereby extending lifespan via a mechanism distinct from rapamycin inhibition that fails to reduce 4E-BP phosphorylation *in vivo*. Finally, Lsp2 adipokine promotes 4E-BP phosphorylation and acts systemically across tissues to shape the lifespan responses to dietary protein. Collectively, our findings establish Lsp2 as a novel translational regulator of TOP genes that mechanistically couples physiological ribosomal protein synthesis with organismal longevity.

## Introduction

Reducing the activity of nutrient signaling pathways via dietary restriction (DR), particularly of amino acids, ameliorates ageing and ageing-associated diseases ^5–7^. A central mediator of dietary restriction in countering ageing is the mechanistic target of rapamycin (mTOR) pathway. The mTOR kinase assembles into two protein complexes, mTORC1 and mTORC2, which respond to distinct environmental and physiological cues and phosphorylate different substrates to adjust organismal metabolism ^1^. mTORC1 senses mainly nutrient availability to tip the balance between anabolism and catabolism in the cell. By phosphorylating the eukaryotic initiation factor 4E-binding protein (4E-BP) and p70 S6 kinase (S6K), mTORC1 is central to promoting protein synthesis, arguably the most energy-intensive process that is tightly linked to organismal health and lifespan ^1,8,9^. Genetically or chemically downregulating the mTORC1 pathway extends lifespan in diverse organisms ^1,10,11^. Particularly, rapamycin, a potent chemical inhibitor of mTORC1 that exerts robust geroprotective effects in mammals, has spurred substantial interests from anti-ageing research ^12^. However, chronic rapamycin treatment causes negative effects on metabolism and immunity, due to its concurrent inhibition of mTORC2 that regulates mainly glucose metabolism ^12^. Thus, there is still intense interest in identifying new ways to specifically inhibit mTORC1 activation and its downstream outputs.

While mTORC1 translationally regulates all mRNAs, it exerts its most rapid and pronounced effect on mRNAs encoding proteins of the translation machinery including all the 79 ribosomal proteins (RPs). These ∼100 mRNAs referred to as TOP mRNAs share a 5′-terminal oligopyrimidine motif (TOP) directly adjacent to the 5′ cap and collectively comprise 15–20% of total cell transcripts ^2^. It has been a long-standing mystery how the presence of a TOP motif on an mRNA renders its the translation exquisitely sensitive to mTORC1-mediated sensing of nutrients ^2,13^. Consistently, varying nutritional conditions ^14^ and ageing ^15^ result in alterations in ribosome composition and levels. Interestingly, TOP motifs are found in RPs of *Drosophila* and vertebrates but not in lower eukaryotes including yeast and worms ^2,16^, implying this mechanism is a recently evolved switch in how nutrient sensing controls growth. Nevertheless, previous studies of TOP motif-mediated translational regulation were done exclusively using cultured cells, thus mechanisms by which the translation of TOP mRNAs responds to nutrients remain entirely unexplored at the organismal level. Considering the significant energy cost in making RPs ^9,17^, identifying *in vivo* mechanisms controlling physiological changes in TOP mRNA translation is expected to open new avenues to target ageing.

Key regulators of TOP mRNA include 4E-BPs ^3,18^, evolutionarily conserved translational repressors that adjust the levels of 5′ cap-dependent mRNA translation according to the nutritional and physiological status of the cell ^1,19,20^. In unphosphorylated state, 4E-BPs suppress translation by binding and sequestering the cap-binding protein eIF4E, thereby preventing its association with eIF4G to form the eIF4F initiation complex. Upon phosphorylation by mTORC1 and potentially other kinases, 4E-BPs release eIF4E, allowing translation initiation ^21,22^. Of note, the allosteric mTORC1 inhibitor rapamycin was found ineffective in slowing down protein translation ^23–25^. In fact, mTORC1-dependent but rapamycin-resistant phosphorylation of 4E-BP instructs key processes including TOP mRNA translation ^3,26,27^. Moreover, forced overexpression of 4E-BP was shown to provide health benefits in flies and mammals ^28–30^. As 4E-BP function is physiologically modulated, this suggests an opportunity to optimize the levels of TOP mRNA translation for lifespan extension by rationally targeting 4E-BP activity.

The fly model has been instrumental in ageing research ^6,31^. *Drosophila* encodes a single 4E-BP by the *Thor* gene that has roles in energy homeostasis and stress responses ^21,28,32–34^. Here we uncover a nutrient-induced adipokine that regulates lifespan by controlling 4E-BP-dependent RP translation. Intriguingly, suppressing nutrient-induced RP translation robustly increases longevity without compromising normal physiology, stress resistance and reproduction of the animal. Our study thus suggests the promising concept that physiological changes in RP translation could be targeted for safe and effective lifespan extension.

## Results

### Adipokine Lsp2 is induced by the “EAA - Insulin - mTORC1” axis

Dietary restriction extends lifespan largely due to the restriction of essential amino acids (EAA)^35^. To uncover physiological alterations incurred by protein diets and during ageing, transcriptomic analyses were performed on the heads of adult flies treated either with a control diet (S group: 2% sucrose) or a protein-rich diet (YE group: 2% sucrose + 3.3% yeast extract) and flies of different age raised on a regular lab diet (Young group: 3 days; Old group: 50 days). Of note, the fly head harbors parts of the fat body, an organ combining metabolic features of mammalian liver and adipose cells ^36–38^. Principal component analysis (PCA) showed that protein ingestion and ageing similarly shifted the gene expression profiles, resulting in a large overlap in the co-regulated genes (Figs. 1a,b and Extended Data Fig. 1a). This included *Lsp2* (Figs. 1c,d and Extended Data Fig. 1b), which encodes a larval serum protein (LSP) specifically expressed in the fat body of feeding larvae in response to AAs and mTORC1 signaling ^4,39^. During development, LSPs are secreted into the haemolymph as storage proteins to sustain the progression of subsequent non-feeding stages. We confirmed the strong induction of *Lsp2* by protein diets in the adult with quantitative PCR (qPCR) and Western blot (Fig. 1e and Extended Data Fig. 1e). In contrast, the remaining LSP genes (*Lsp1α*, *β* and *γ*) were neither expressed (FPKM<0.1) nor induced by protein diets (Fig. 1e and Extended Data Figs. 1c,d), hinting at an additional role of Lsp2 in adult physiology beyond its storage function. *Lsp2* was predominantly expressed in the head fat body and its induction was independent of sex and reproductive status of flies (Extended Data Figs. 1f-h).

**Fig. 1.**
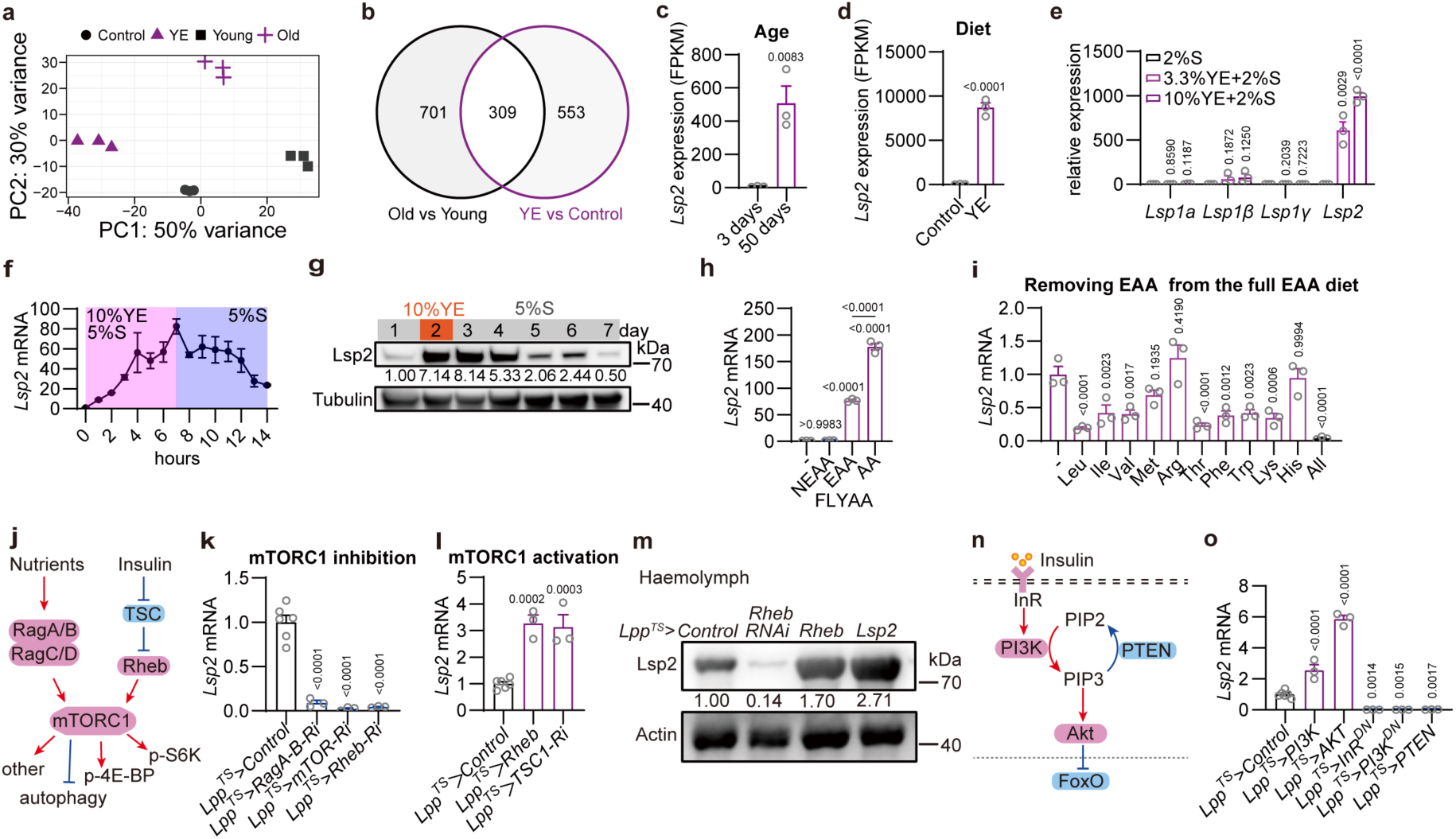
Induction of Lsp2 from the fat body by essential amino acids requires mTORC1. **a-d**, RNA-seq reveals ageing and protein diets induce overlapping transcriptional changes including the induction of *Lsp2*. PCA analysis (**a**) of differentially expressed genes (DEGs, *Q*<0.05 & |log_2_FoldChange|>1) in heads of adult females ingesting different diets (Control, 2% sucrose; YE, 2% sucrose with 3.3% yeast extract (YE); 5-day-old *w*^111^*^8^* flies were kept for 2 days on Control or YE diets) or of different ages (Young, 3-day-old flies; Old, 50-day-old flies; *Oregon R* flies raised on our conventional lab diet (CD)), and comparison of DEGs from the two datasets (**b**) with *Lsp2* expression shown (**c**,**d**). **e-g**, *Lsp2* but not other *LSPs* is reversibly induced by protein diets. qPCR shows relative expression of the indicated *LSP* genes in heads of 5-day-old *w^1118^* flies ingesting Control (2%S) or YE-based protein diets for 2 days (**e**). Dynamic changes in *Lsp2* transcript (**f**, using fly heads) and Lsp2 protein (**g**, using whole flies) levels in 5-day-old flies that were first on 5% sucrose for 1 day and then on a protein diet (10%YE+5%S) for 7 hours before returning to 5% sucrose for another 7 hours. **h**,**i**, *Lsp2* is induced by diets with balanced EAAs. qPCR detecting relative expression of *Lsp2* in heads of 5-day-old flies ingesting a mixture of 20 AAs, 10 EAAs or 10 NEAAs, respectively (**h**), or a full EAA diet lacking only one indicated EAA (**i**) for 1 day. AAs were provided according the FLYAA recipe with 1% sucrose added. **j-m**, Lsp2 is controlled by mTORC1 signaling and released from fat body to haemolymph. A schematic of mTORC1 signaling (**j**); relative *Lsp2* levels in heads of flies with the indicated genotypes to inhibit (**k**) and activate (**l**) mTORC1 signaling in the fat body using *Lpp^TS^* as driver; and circulating Lsp2 protein levels in haemolymph of flies with the indicated genotypes (**m**). Flies were kept on CD for 5 days at 29℃ to induce transgene expression. **n**,**o**, *Lsp2* is controlled by Insulin signaling. A schematic of Insulin signaling (**n**) and relative *Lsp2* levels (**o**) in heads of flies with the indicated genotypes kept on CD for 5 days at 29℃. 20 females were used for each sample and represented as one dot except **f**. Data are mean ± s.e.m. Student’s *t*-test in **c**,**d**; one-way ANOVA with Dunnett’s multiple comparisons test in **e**,**h**,**i**,**k**,**l**,**o**. *P* values are indicated.

Importantly, the transcriptional induction of *Lsp2* by protein diet was reversible, increasing for about 80 times within 7 hours but lasting only several hours upon withdrawal of the protein source (Fig. 1f). This raised the possibility that *Lsp2* transcript levels duly report the fluctuations in systemic protein nutrition of the organism. Lsp2 protein instead appeared more stable and lasted about 2 days after Lsp2 induction by a protein diet (Fig. 1g). As protein diets of other sources (Tryptone and BSA) also increased *Lsp2* expression (Extended Data Figs. 1i,j), we further tested *Lsp2* induction by amino acids. A mixture of the 10 EAAs rather than 10 non-essential amino acids (NEAA) ^40^ strongly induced *Lsp2*, although NEAAs further increased *Lsp2* expression only in the presence of full EAAs (Fig. 1h). *Lsp2* expression appeared to reflect a balance in systemic EAA levels, as removing most individual AA from the full EAA diet significantly reduced *Lsp2* induction (Fig. 1i), although methionine, histidine and arginine appeared unnecessary, reminiscent of a recent report ^41^ and the absence of arginine sensors in *Drosophila* ^1^. Moreover, single EAA provided at sufficient amount only marginally induced *Lsp2* (Extended Data Fig. 1k). Additional tests of a few AA combinations suggested it unlikely that individual or a subset of EAAs can fully induce *Lsp2* (Extended Data Figs. 1l,m), placing *Lsp2* levels as a *bona fide* indicator of balanced EAAs in the diet.

As multiple EAAs activate mTORC1 ^1^, we tested if adult *Lsp2* expression is dependent on mTORC1 signaling (Fig. 1j). A fat body-specific *Lpp-Gal4* combined with *tub-Gal80^TS^* (referred as *Lpp^TS^>*) was used for adult-onset transgene expression. Inhibiting mTORC1 signaling by knocking down *RagA-B*, *mTOR* or the *Rheb* GTPase all significantly reduced *Lsp2* (Fig. 1k), while increasing mTORC1 activation by knocking down *TSC1* or overexpressing *Rheb* both increased *Lsp2* expression (Fig. 1l). Genetic manipulation of mTORC1 activity in the fat body changed the levels of circulating Lsp2 protein in fly haemolymph (Fig. 1m), validating Lsp2 as an mTORC1-induced circulating adipokine. In addition, Insulin/IGF signaling, an evolutionarily conserved regulator of animal lifespan ^42,43^, also activates mTORC1 ^44^. Activating Insulin signaling by expressing PI3K or AKT significantly upregulated *Lsp*2, while suppressing it via overexpressing PTEN or dominant negative forms of insulin receptor (*InR^DN^*) and PI3K (*PI3K^DN^*) all reduced *Lsp2* levels (Figs. 1n,o). Thus, the Insulin-mTORC1 nutrient-sensing axis controls the expression of Lsp2 that appears to decode a balance in systemic EAA levels.

### Robust lifespan extension in *Lsp2* mutants

We wondered if Lsp2 might act as an adipose effector of mTORC1 in modulating the organismal physiological response to nutrients. As mTORC1 signaling has a prominent role in lifespan ^10,11,45,46^, we generated *Lsp2* mutants and checked their lifespan. Mobilization of a transposon in the *Lsp2* locus produced both a revertant control *Lsp2^B^*^14^ via precise excision and an insertional mutant *Lsp2^A^*^10^ that showed diminished *Lsp2* transcripts and lacked detectable Lsp2 protein (Extended Data Figs. 2a-c). As *Lsp2^B14^* and *Lsp2^A10^* lines share the same genetic background, their lifespan is readily comparable. An *Lsp2* knockout (*Lsp2^KO^*) with most of the *Lsp2* coding sequences removed was also generated using CRISPR/Cas9 ^47^. Both *Lsp2^A10^*and *Lsp2^KO^* flies were homozygous viable and fertile with no apparent defects. In addition, *Lsp2^A10^* mutants had no developmental delay in pupation and adult eclosion except that *Lsp2^A10^* larvae pupated slightly faster (Extended Data Figs. 2d,e), in line with a recent report ^4^.

Then, the lifespan response to various dietary protein levels was measured using the sugar-yeast-agar (SYA) diets. These diets all contained 5% sucrose but varied in the content of yeast as protein source ranging from 0-20% (w/v). As expected, Lsp2 protein levels exhibited a dose-dependent increase in response to elevated dietary protein concentrations (Fig. 2a). Consistent with a previous report ^46^, either strong limitation or excess of dietary yeast were not optimal for lifespan, and SYA foods containing 5% yeast (0.5X SYA foods) maximized median lifespan of control females (Figs. 2b-e and Extended Data Fig. 2f). Interestingly, *Lsp2^A10^*females showed increased longevity compared to controls on all the SYA diets tested, with median lifespan increased by 14%, 18%, 28% and 17%, on 0.1X, 0.5X, 1.0X and 2.0X SYA food, respectively (Figs. 2b,c). *Lsp2^KO^* females had similar lifespan responses (Extended Data Fig. 2f). Of note, both *Lsp2* mutants tolerated well the negative effects of protein-rich diets (1.0X and 2.0X SYA foods) on lifespan. Conversely, overexpressing *Lsp2* consistently shortened lifespan, with median lifespan reduced by 38%, 37%, 20%, 20% and 4%, on 0.0X, 0.1X, 0.5X, 1.0X and 2.0X SYA food, respectively (Figs. 2d.e). The more pronounced lifespan reduction of females overexpressing *Lsp2* during protein scarcity (0.0X and 0.1X SYA foods) implied that Lsp2 promoted an energy-intensive cellular process. Cox proportional hazard analysis confirmed that the interaction between genotypes and dietary protein levels was both significantly changed in *Lsp2^A10^* flies (*P* = 0.0404) and *Lpp^TS^>Lsp2* flies (*P* = 0.0006). Notably, this is reminiscent of lifespan regulation by altering mTORC1 activity ^46^, indicating that Lsp2 is an adipose effector of mTORC1 signaling in mediating lifespan response to dietary protein.

**Fig. 2.**
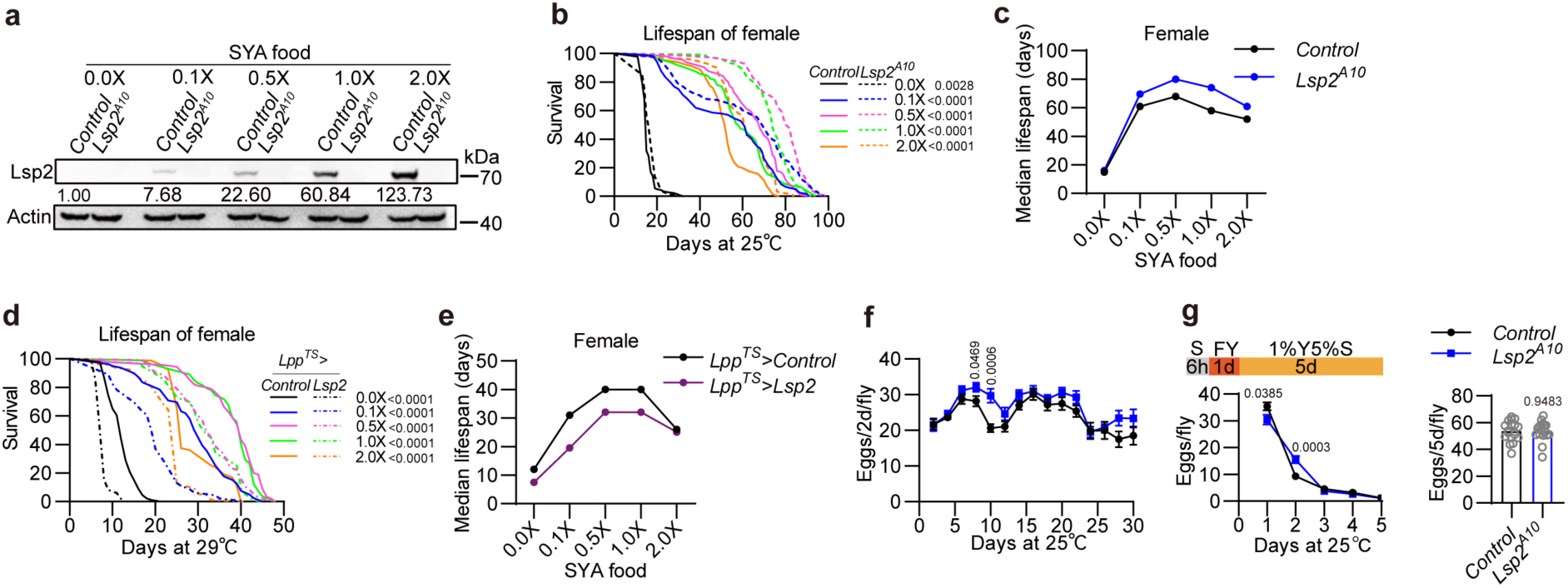
Lifespan extension in *Lsp2* mutants without costs in reproduction. **a**, Lsp2 protein levels in *Control* and *Lsp2^A1^*^0^ mutant flies raised on SYA (sugar-yeast-agar) foods of increasing yeast contents (0%, 1%, 5%, 10% and 20% as 0.0X, 0.1X, 0.5X, 1.0X and 2.0X SYA diets, respectively) for 14 days. Whole flies were used. **b-d**, Lsp2 negatively regulates lifespan. Survival (**b**,**d**) and median lifespan (**c**,**e**) of flies with the indicated genotypes on the five SYA diets. For *Lsp2^A1^*^0^ mutants, n = 127, 126, 114, 110 and 122; for its genetic control, n = 141, 121, 113, 118 and 126 (**b**,**c**). For *Lsp2* overexpression from the fat body, n = 98, 46, 95, 96 and 93; for its control, n = 86, 79, 90, 82 and 83 (**d**,**e**). **f**,**g**, Lsp2 is required for timely reproduction but not for fecundity. Monitoring the egg laying of a group of 3 females with 3 males (1-day old) every two days for 30 days for *Control* (n = 15) and *Lsp2^A10^*(n = 17) flies raised on conventional lab diet (**f**). Egg laying of *Control* (n = 17) and *Lsp2^A10^* (n = 17) flies during a fasting-refeeding treatment (**g**), where a group of 5 females with 3 males (5-day old) were dry-starved for 6 h and allowed to ingest fresh yeast (FY) for one day before being shifted to the 0.1X SYA diet. Shown are eggs laid each day (*left*) and over five days (*right*). Data are mean ± s.e.m. in **f**,**g** with Student’s *t*-test. Log-rank test in **b**,**d**. *P* values are indicated.

Essentially similar lifespan responses were also observed for male *Lsp2^A10^* and *Lpp^TS^>Lsp2* flies, although sex differences did exist (Extended Data Figs. 2g,h). This is in line with the fact that mated males and females consume proteins largely for different purposes and differ in protein requirements to maximize lifespan ^34,48,49^. In addition, loss of *Lsp2* also extended the lifespan on conventional lab diets (CD) (Extended Data Figs. 2i,j). Overexpressing *Lsp2* shortened lifespan on both CD and a diet using yeast extract as an alternative protein source, and the lifespan shortening was also seen in *Lsp2* overexpression using additional Gal4 drivers and a *UAS-Lsp2* transgene (Extended Data Figs. 2k-p). Reduced proliferative activity in ageing intestinal epithelia generally correlates with the increase in longevity ^50,51^. Agreeing with this notion, loss of *Lsp2* decreased intestinal stem cell (ISC) proliferation while *Lsp2* overexpression promoted it (Extended Data Figs. 2q,r). Taken together, Lsp2 molecularly connects protein nutrition to lifespan regulation.

### No major life-history cost is associated with lifespan extension in *Lsp2* mutant

Decreased fecundity and increased stress resistance are often associated with lifespan extension ^35,52–54^. However, the long-lived *Lsp2^A^*^10^ flies were largely spared these tradeoffs. First, this mutant did not show altered resistance either to starvation or to oxidative stress (Extended Data Figs. 3a,b). Second, we detected no difference in the number of eggs laid, climbing ability, spontaneous locomotor activity, and survival to systemic bacterial infection in both 10-day-old and 30-day-old flies (Extended Data Figs. 3c,d). Third, monitoring egg laying consecutively for 30 days did not reveal any decline in the fecundity of *Lsp2^A10^* flies compared to control (Fig. 2f). In fact, 8-10-day-old *Lsp2* mutants exhibited significantly increased egg laying, indicating that lifespan extension in *Lsp2* mutants unexpectedly came with improved reproduction. Finally, genetic manipulation of *Lsp2* expression also impacted the lifespan of male flies, further arguing against a lifespan and fecundity tradeoff. Thus, lifespan extension in *Lsp2* mutants did not seem to associate with major costs in life-history traits.

As *Lsp2* is strongly but transiently induced by dietary protein that is vital for insect reproduction ^55^, we tested if Lsp2 promoted timely reproduction. Under a starvation-refeeding cycle, *Lsp2^A10^*flies laid significantly fewer eggs than control flies on the first day of yeast refeeding (Fig. 2g), although the total number of eggs laid over the five days after yeast refeeding did not differ. This indicated that Lsp2 is required to promptly adapt fly reproduction to protein-rich food that might appear limited in the field.

### Lsp2 is a translational regulator that selectively promotes TOP mRNA translation

To dissect the molecular basis of lifespan regulation by Lsp2, we first performed RNA-seq analyses to detect transcriptomic changes. While overexpressing *Rheb* to activate mTORC1 signaling changed the expression of 684 genes, *Lsp2* overexpression in sharp contrast altered only 35 genes (Extended Data Figs. 4a-e). This primarily excluded Lsp2 as a transcriptional regulator. Nevertheless, *hsp70Ba* and *hsp70Bb*, two Hsp70 genes encoding a molecular chaperone for folding of newly synthesized proteins thereby maintaining translational homeostasis ^56^, were among the 35 differentially expressed genes (Extended Data Figs. 4b,f). qPCR showed that expressing *Lsp2* from the fat body upregulated *Hsp70Ba* mRNA in several tissues, indicating Lsp2 exerts a systemic effect (Extended Data Fig. 4g). We thus suspected that upregulation of *Hsp70* factors by Lsp2 stemmed from protein folding stress associated with elevated translation, an energy-demanding process ^20,57^.

Levels of protein translation are dynamically tailored to environmental and physiological stresses sensed by specialized kinases that converge on phosphorylation of the α subunit of eukaryotic initiation factor 2 (p-eIF2α) that inhibits global translation ^58^. We found that loss of *Lsp2* increased while *Lsp2* overexpression reduced p-eIF2α levels in whole flies raised on SYA diets of various protein concentrations (Extended Data Fig. 4h). Interestingly, the change in p-eIF2α levels occurred both locally in the fat body and remotely in the gut (Extended Data Fig. 4i), reminiscent of the systemic *Hsp70* upregulation by Lsp2. Consistently, puromycin incorporation assay revealed reduced global protein synthesis in *Lsp2* mutant flies (Extended Data Fig. 4j). Thus, Lsp2 is required for translation. Reducing protein translation extends lifespan in various organisms ^8,59–61^, and likely underlies the increase in longevity of *Lsp2* mutant flies.

Translatomics and proteomics were then performed to reveal how Lsp2 impacts translation *in vivo*. Whole animal lysates of equal RNA content were fractionated via sucrose density gradient centrifugation, followed by quantification of UV absorbance of each fraction. The polysome profiling analyses indicated that the amount of polysome-bound RNA in *Lsp2^A10^* flies tended to decrease (Fig. 3a). Of note, a reduction in the 40S small subunit but increase in the 60S large subunit simultaneously occurred in *Lsp2* mutants (Figs. 3b,c). Such an imbalance between ribosomal subunits potentially impairs ribosome assembly, thereby limiting translation ^62,63^. Next, monosome-bound and polysome-bound mRNAs were processed for deep sequencing (monosome-seq and polysome-seq), along with regular transcriptomics to assess the overall abundance of mRNAs. At the transcriptomic level, only 33 genes were upregulated, while 305 genes were downregulated with a significant enrichment in the Toll and Imd signaling pathways. To identify mRNAs significantly altered by Lsp2 at the translational level, we calculated the translational efficiency (TE) of each mRNA by normalizing their expression in the translatome to the corresponding transcript (Fig. 3d). This detected 66 TE-downregulated but only 6 TE-upregulated mRNAs in *Lsp2* mutants (Extended Data Table 1). Functional clustering revealed pronounced enrichment of mRNAs coding for RPs (Fig. 3e and Extended Data Figs. 5b,c). Strikingly, 48 out of 66 TE-downregulated genes were cytosolic RPs (Extended Data Fig. 5a). In fact, nearly all the other cytosolic RPs tended to be translationally suppressed by the loss of *Lsp2* (Fig. 3d). In sharp contrast, translation of mitochondrial RPs was not significant altered (Extended Data Figs. 5h; 6a,b). This indicates that Lsp2 is selectively required for global translation of genes coding for cytosolic RPs. In addition, overexpressing *Lsp2* or *Rheb* that robustly induces *Lsp2* both broadly upregulated the translation of RP genes revealed again by polysome profiling followed by deep RNA sequencing (Extended Data Figs. 7a,b).

**Fig. 3.**
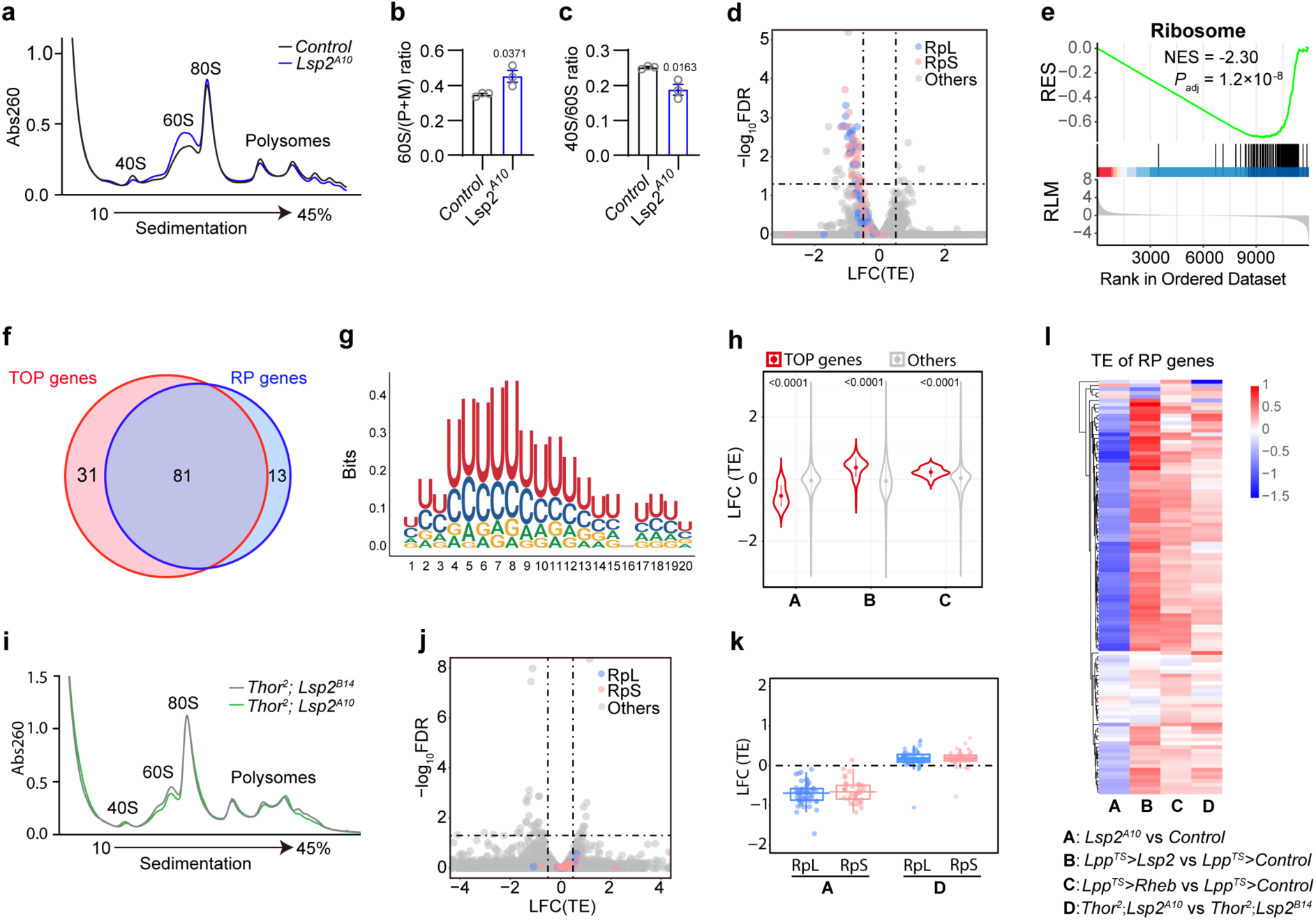
Lsp2 selectively promotes TOP mRNA translation through 4E-BP. **a-e**, Lsp2 is required for global RP translation. Polysome profiling (**a**) of 15-day-old *Control* and *Lsp2^A10^*mutant flies raised on 0.5X SYA food reveals an increase in the 60S subunit (**b**) but a decrease in the 40S subunit (**c**) in *Lsp2* mutants. A volcano plot of translational efficiency (TE) changes in *Lsp2* mutants with RP mRNAs highlighted (**d**) reveals a global downregulation of RP translation. The dashed lines denote thresholds. Gene Set Enrichment Analysis (GSEA) indicates significant downregulation of the Ribosome pathway in *Lsp2* mutants (**e**). **f-h**, Features of the predicted fly TOP genes. Overlap between fly TOP genes and RP genes (**f**); sequence logo of fly TOP motif (**g**) generated from the 5′ end of RP mRNAs; and changes in the TE of fly TOP genes (**h**) in conditions of *Lsp2* loss (group A) and *Lsp2* gain of function (groups B and C). **i-l**, Loss of *4E-BP* (fly *Thor*) suppresses the reduction in RP translation seen in *Lsp*2 mutants. Polysome profiling (**i**) of 15-day-old *Thor^2^; Lsp2^B14^* and *Thor^2^; Lsp2^A10^* flies raised on 0.5X SYA food, and a volcano plot (**j**) showing the associated TE changes with RP mRNAs highlighted. Further shown are a summary of changes in TE of RP genes in experimental groups A and D in a box and whisker plot (**k**), and a heatmap of changes in TE of RP genes across the four experimental groups (**l**). Data are presented as mean ± s.e.m. in **b**,**c** with Student’s *t*-test. Permutation test in **e**. Wilcoxon test in **h**. *P* values are indicated.

Despite a strong bias in translation of RP genes, proteomic analysis revealed no significant changes in the overall abundance of RPs, though a tendency to decrease globally was observed (Extended Data Fig. 5h). This was corroborated by Western blots to assess the total amounts of several RPs (Extended Data Fig. 5i). It is possible that post-translational protein quality control mechanisms are at work to adjust global RP levels ^64,65^, although polysome profiling did suggest a difference between *Lsp2* mutant and control flies in RPs assembled into ribosomal subunits (Fig. 3a).

RPs constitute a major part of the TOP mRNAs that are remarkably abundant in cells ^2^. Using a published algorithm ^16^, we calculated TOP scores for each *Drosophila* mRNAs and identified in total 112 TOP genes (with TOP score >3). Like mammalian TOP mRNAs, the predicted fly TOP mRNAs also predominantly encode RPs (81 out of 112; *Drosophila* encodes in total 94 RP genes, making 79 RPs) and other translational regulators (Fig. 3f and Extended Data Table 2). Compared with the canonical TOP motifs present in mammalian RP genes, fly RP genes appear to feature a distinct oligopyrimidine motif (referred to as fly TOP motif) that less frequently starts with a cytidine (C) while consisting of longer uniform stretch of U/C rich sequences (Fig.3g and Extended Data Figs. 8a,b). To reveal if the predicted fly TOP motif is functionally relevant, we closely looked at the 5′ regions immediately after the transcriptional start site (TSS) in mRNAs of the 66 genes translationally downregulated due to the absence of the mTORC1 effector Lsp2. With mRNA isoforms also counted for this analysis, we found that, out of the 48 *Drosophila* RP genes, only 11 harbor canonical TOP motifs that are found in all human RP genes ^66^. Moreover, another 11 possess the *iso-TOP* motif defined as sequences otherwise identical to canonical TOP motifs except for a substitution of cytidine (C) to uridine (U) at TSS, while the remaining 26 RPs feature extended pyrimidine-rich sequences within the first 50 nucleotides of their mRNAs (Extended Data Table 3). Such sequence flexibility in fly TOP motifs is in line with the notion raised previously that TOP motifs can tolerate certain degrees of sequence diversification while remaining responsive to translational control ^3^. In contrast, among the 18 translationally downregulated non-RP genes, only 5 carry pyrimidine-rich sequences, and none shows clear canonical TOP or iso-TOP motifs, further hinting at the mechanistic complexity of mRNP-mediated translational control ^13,66^.

As RP mRNAs of worm, yeast, *Arabidopsis* and rice all lack TOP and similar motifs (namely iso-TOP and pyrimidine-rich motifs) (Extended Data Fig. 8a), we speculate *Drosophila* iso-TOP motifs and the pyrimidine-rich sequences likely represent the primitive forms of mammalian TOP motifs. These may have acquired the ability to mediate a translational response to nutrients at least via the action of the Lsp2 adipokine, thereby placing the translation apparatus under nutrient-dependent translational control. Finally, a re-analysis of our translatomic data indicated that these predicted fly TOP genes are indeed selectively modulated by both the loss and gain of Lsp2 function (Fig. 3h). Together, these findings reveal that Lsp2 is a specific translational regulator of mRNAs with TOP and similar motifs.

### Lsp2 regulates TOP mRNA translation and lifespan through 4E-BP

TOP mRNAs are translationally regulated by the mTORC1 pathway ^3,18^. Of note, complete mTORC1 inhibition with ATP site mTOR inhibitors rather than with rapamycin reduces the translation of TOP mRNAs ^3^. As knockout of 4E-BPs renders TOP mRNA translation resistant to mTORC1 inhibition in mammalian cells, we wondered if the reduction in global RP translation as seen here in the *Lsp2* mutant could be similarly suppressed by loss of *4E-BP* (encoded by *Thor* in *Drosophila*). To this aim, both the *Lsp2^A10^* mutant and *Lsp2^B14^*control chromosomes were crossed to a null allele of *4E-BP* (*Thor^2^*), and the *Thor^2^*; *Lsp2^A10^* and *Thor^2^*; *Lsp2^B14^* flies were subjected to further translatomic analysis (Fig. 3i). Ribosome profiling revealed that the imbalance in the 40S and 60S subunits observed in *Lsp2* mutant flies was now abolished (Extended Data Fig. 5j). In addition, global protein translation in the *Lsp2* mutant measured by the puromycin incorporation assay was also reverted to control levels in the *Thor^2^* background (Extended Data Fig. 5d). We next performed monosome-seq and polysome-seq along with matched mRNA-seq to detect translational changes (Extended Data Figs. 5e,f). Strikingly, the global reduction in RP translation seen in the *Lsp2* mutant was also completely suppressed by the loss of *4E-BP*, although *Thor^2^; Lsp2^A10^* and *Thor^2^; Lsp2^B14^* flies still differed in the translational levels of many other genes (Figs. 3j-l and Extended Data Fig. 5k). Thus, the adipokine Lsp2 promotes RP translation genetically via 4E-BP. To further assess the role of RP translation in lifespan, we compared the longevity of *Thor^2^; Lsp2^A10^* flies to *Thor^2^; Lsp2^B14^* control flies raised on CD and on SYA diets of different protein levels. Intriguingly, lifespan extension in the *Lsp2* mutant was largely abolished by simultaneously removing *4E-BP* (Figs. 4a,b and Extended Data Figs. 5g,l). These findings placed 4E-BP as the crucial mediator of Lsp2 functions in RP translation and lifespan.

**Fig. 4.**
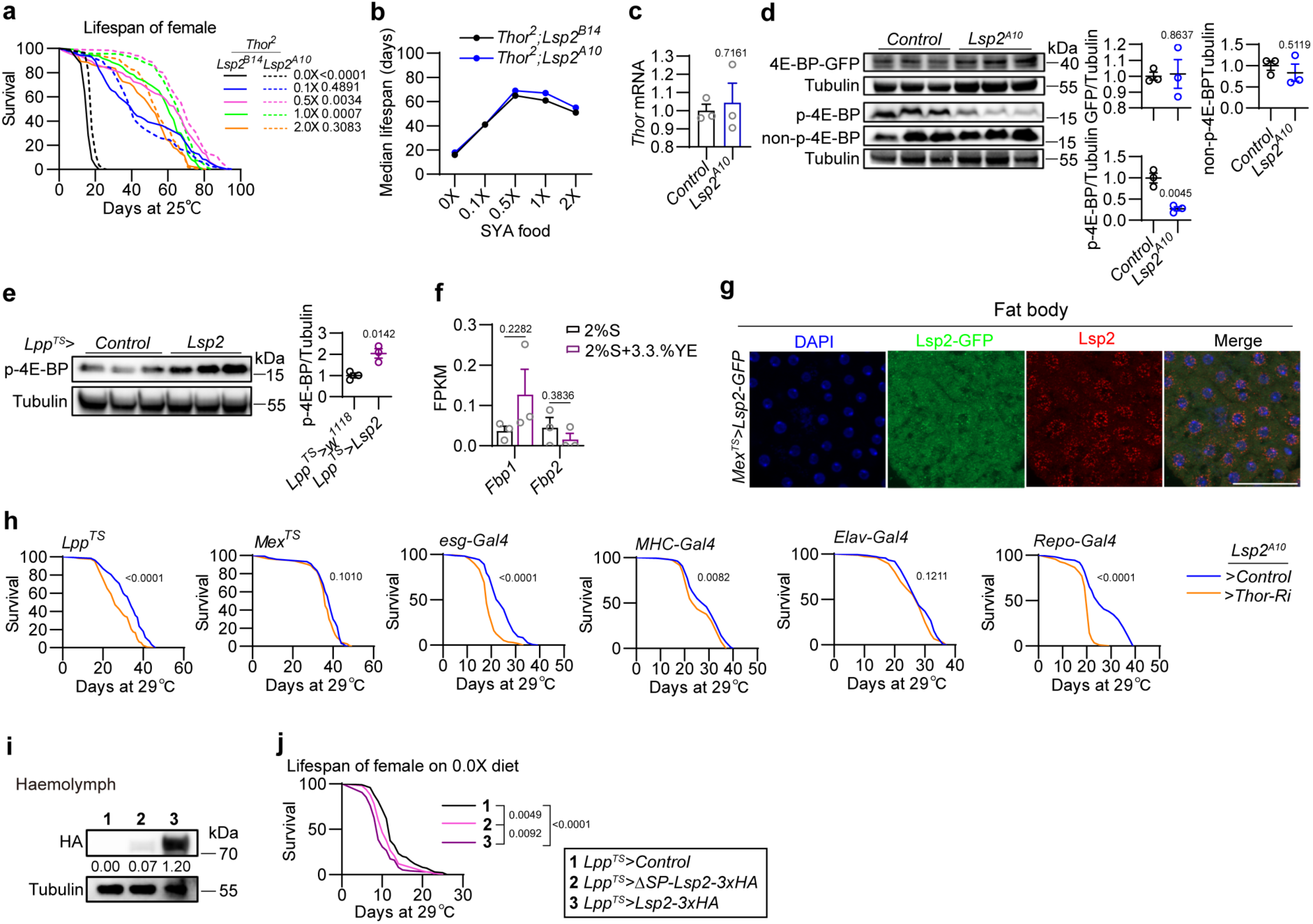
Loss of *Lsp2* extends lifespan by reducing 4E-BP phosphorylation. **a**,**b**, Loss of *4E-BP* abolishes the lifespan extension in *Lsp2* mutants. Survival (**a**) and median lifespan (**b**) of flies with the indicated genotypes on the five SYA diets. For *Thor^2^; Lsp2^A10^*, n = 114, 114, 126, 129 and 113; for *Thor^2^; Lsp2^B14^*, n = 120, 127, 113, 97 and 121. **c-e**, Lsp2 controls 4E-BP phosphorylation. qPCR detecting relative *Thor* mRNA levels (**c**) and Western blots (**d**,**e**) measuring total 4E-BP protein (using a *Thor-GFP* protein trap), Thr46 dephosphorylated form (non-p-4E-BP) or Thr37/46 phosphorylated 4E-BP (p-4E-BP) levels in flies with the indicated genotypes. Samples used are heads of 7-day-old flies raised on conventional lab diet (CD) (**c**), 7-day-old whole flies raised on CD (**d**), and whole flies raised at 29℃ on CD for 3 days and then on 5% sucrose for 3 days (**e**). One dot in quantification graphs stands for 20 flies. **f**,**g**, Two putative Lsp2 receptors *Fbp1* and *Fbp2* are neither expressed in adults nor induced by protein diets (**f**, data from RNA-seq described in Fig. 1a), but Lsp2-GFP expressed from the gut can still be internalized into the fat body cells in adults (**g**). Flies in **g** were raised at 29℃ on CD for 5 days. **h**, Requirements of *4E-BP* in multiple tissues for the lifespan of *Lsp2^A10^* flies. Survival analyses of *Lsp2* mutants expressing an RNAi *Control* or a *Thor* dsRNA using *Lpp-Gal4^TS^* (fat body, n = 140 and 127), *Mex-Gal4^TS^* (enterocytes, n = 121 and 116), *esg-Gal4* (intestinal stem cells, n = 157 and 168), *MHC-Gal4* (muscles, n = 161 and 133), *Elav-Gal4* (neurons, n = 136 and 76) and *Repo-Gal4* (glial cells, n = 162 and 165), performed at 29℃ on CD. Females were used. **i**,**j**, Expressing an Lsp2 variant that lacks the signal peptide (ΔSP-Lsp2-3xHA) still shortens fly lifespan. Western blots (**i**) reveal that ΔSP-Lsp2-3xHA expressed in the fat body was barely detected in the haemolymph of flies raised at 29℃ on CD for 7 days. 50 females/sample were used to collect haemolymph. Note that expressing *ΔSP-Lsp2-3xHA* in the fat body shortened lifespan but to a lesser extent compared to the effect of *Lsp2-3xHA* (**j**). Lifespan was monitored on 0.0X SYA food. n = 99, 101 and 75 for flies of the three indicated genotypes. Data are mean ± s.e.m. in **c-f**. Log-rank test in **a**,**h**,**j**; Student’s *t*-test in **c-f**. *P* values are indicated. Scale bar 50 µm in **g**.

We noticed that, besides RP translation, Lsp2 concurrently increased mTORC1 activity thereby forming a feedback amplification loop to sustain its robust upregulation by EAAs (Extended Data Figs. 9a-c), and food intake via remotely modulating insulin secretion (Extended Data Figs. 9d-k). Nevertheless, removing *4E-BP* failed to suppress the decreases in mTORC1 activity and food intake seen in *Lsp2* mutant (Extended Data Figs. 9l,m), suggesting that these two aspects did not occur through 4E-BP and thus might not significantly contribute to the lifespan extension in *Lsp2^A10^* flies. Furthermore, the reduction in ISC proliferation seen in aged *Lsp2^A10^* flies was still maintained in the *4E-BP* null background (Extended Data Fig. 9n). We therefore conclude that loss of *Lsp2* extends lifespan primarily by restricting RP translation.

Next, we addressed how Lsp2 modulates 4E-BP. Loss of *Lsp2* did not alter *4E-BP* levels in mRNA, total protein (using a *Thor-GFP* protein trap) and the dephosphorylated form (dephosphorylated at Thr46) (Figs. 4c,d). Nevertheless, Thr37/46 phosphorylated 4E-BP that frees eIF4E to initiate translation was significantly reduced in *Lsp2* mutant but increased upon *Lsp2* overexpression (Figs. 4d,e). As mTORC1 is the best-known 4E-BP kinase and Lsp2 also regulates other mTORC1 targets (Extended Data Figs 9a,b), it is likely that Lsp2 promotes 4E-BP phosphorylation most likely via mTORC1, but further work is required to reveal the details. Nevertheless, structural modeling predicted interactions between Lsp2 hexamer and 4E-BP (Extended Data Figs 10a,b).

We sought to further reveal the target tissues that Lsp2 adipokine acts on to impact lifespan, ideally by identifying the receptor of Lsp2. It was reported that LSPs stored in the haemolymph of late fly larvae as hexamerins are later internalized into the fat body via Fbp1 ^4^. However, these putative LSP endocytic receptors Fbp1 and Fbp2 were no longer expressed in adults or induced by protein diets (Fig. 4f). Moreover, Lsp2-GFP ectopically expressed from the midgut was still detected in the cytoplasm of fat body cells (Fig. 4g), suggesting alternative mechanisms of Lsp2 uptake. As loss of *4E-BP* abolished the lifespan extension of *Lsp2* mutant, we alternatively turned to tissue-specific depletion of *4E-BP* in *Lsp2* mutant and inferred the target tissue by checking lifespan. Knocking down *4E-BP* by RNAi in the fat body, the ISCs and glial cells strongly reduced the lifespan of *Lsp2* mutant, while *4E-BP* depletion in muscles had a marginal effect (Fig. 4h). Finally, expressing in the fat body a non-secretable Lsp2 that lacks the signal peptide still shortened lifespan, but to a lesser extent compared to that caused by wild type Lsp2 (Figs. 4i,j). These results align well with the systemic effects that Lsp2 exerted on Hsp70 and p-eIF2α levels, indicating that the Lsp2 adipokine acts on multiple tissues to coordinate whole organismal physiology thereby shaping the lifespan response to protein diets.

### 4E-BP control of RP translation is a novel longevity target

Since the 4E-BP branch downstream of mTORC1 largely resists rapamycin inhibition in cells ^3,26^, our work highlights the physiological modulation of 4E-BP-dependent RP translation as an unexplored avenue to lifespan extension by targeting specific mTORC1 outputs. To enhance the notion that loss of *Lsp2* extends lifespan via a mechanism distinct from that of rapamycin, we directly compared the impact of rapamycin treatment with that of *Lsp2* mutation on lifespan. Rapamycin treatment extended the lifespan of control flies raised on CD and 1.0X but not 2.0X SYA diets as expected but consistently shortened the lifespan of *Lsp2* mutants under these dietary conditions (Figs. 5a-c). This indicates that the physiological state conferred by *Lsp2* loss is already optimized for longevity, particularly under high protein conditions. Intriguingly, in wild type flies following rapamycin treatment Western blots revealed reduced S6K activity *in vivo* as shown previously ^10^, but surprisingly a simultaneous increase in Lsp2 and p-4E-BP (Thr37/46) levels (notably the p-4E-BP/non-p-4E-BP ratio) (Fig. 5d). The finding that rapamycin promoted 4E-BP phosphorylation at the organismal level is entirely unexpected and warrants further study, but we suspect that Lsp2 is key for this. Thus, unlike rapamycin which is unable to reduce 4E-BP phosphorylation *in vivo*, Lsp2 appears an intrinsic modulator of 4E-BP activity. The greater extent of lifespan extension demonstrated by *Lsp2* loss has principally validated the potential of targeting 4E-BP activity for longevity.

**Fig. 5.**
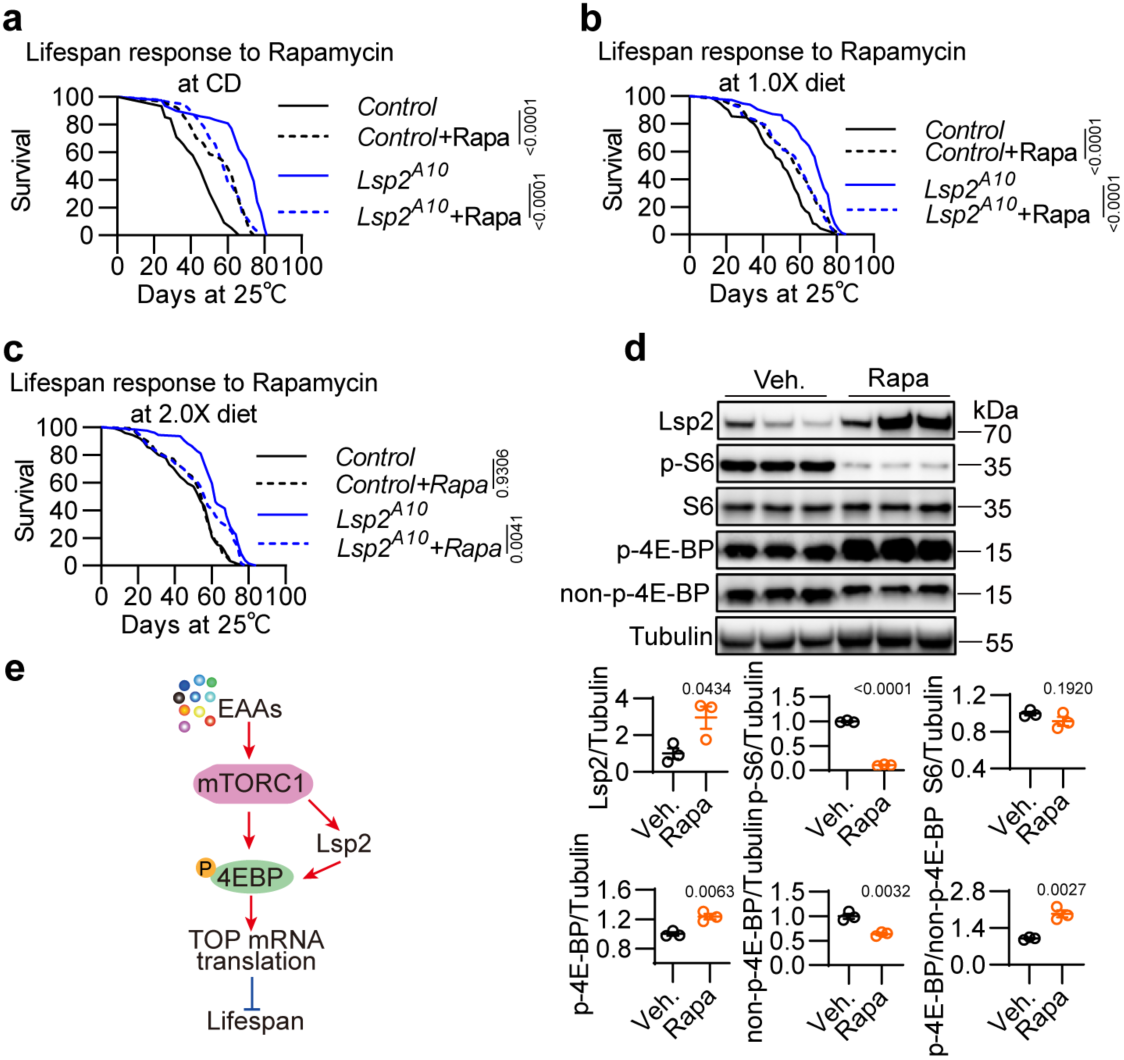
Long-lived *Lsp2* mutants are susceptible to rapamycin treatment. **a-c**, Lifespan analyses of *Control* and *Lsp2^A10^*flies treated with 200 µM rapamycin or its solvent added to conventional lab diet (**a**), 1.0X (**b**) and 2.0X (**c**) SYA foods. **d**, Western blots and quantification of the indicated proteins in 5-day-old *w^1118^*flies treated with the solvent or 200 µM rapamycin in 1.0X SYA food for 10 days. **e**, Model. Flies used for *Control*, *Control*+Rapa, *Lsp2^A10^* and *Lsp2^A10^*+Rapa are 76, 94, 79 and 94 in **a**; 111, 119, 117 and 131 in **b**; 126, 119, 128 and 125 in **c**. Data are mean ± s.e.m. in **d** with Student’s *t*-test. Log-rank test in **a-c**. *P* values are indicated. Veh. for vehicle; Rapa for rapamycin.

## Discussion

A variety of mechanisms are at work to drive animal preference for diets rich in protein ^67–69^. Here, we show that protein diets induce in adipose tissue an mTORC1 effector that acts via 4E-BP to increase physiological RP translation. Strikingly, removing this effector to blunt the translational response to protein diets ensures robust lifespan extension yet without major costs in normal physiology, stress resistance and reproduction to the animal. Given that *Lsp2* is induced only upon protein intake, our study raises the concept that an inducible mechanism is encoded at the organismal level to strengthen cell translation by potentiating mTORC1 outputs. Inhibiting such a physiological “translation-enhancing mechanism” (Fig. 5e), rather than fully turning the nutrient-sensing pathway off, is sufficient to provide health benefits in the context of ageing and overnutrition. While Lsp2 benefits juvenile growth ^4^, our work and an accompanying paper from Obata and colleagues ^70^ both found it shortens adult lifespan by promoting RP synthesis, offering robust evidence for the antagonistic pleiotropy theory of ageing ^71^. By directly connecting the molecular mechanism of a pleiotropic gene to ageing trajectories, our findings advance understanding of how evolutionary forces shape ageing. In an evolutionary perspective, insect hexamerins seem to share functional parallels with immunoglobulin G (IgG), one of the most abundant proteins in human serum that comprises ∼10–20% of plasma proteins ^72^. Notably, IgG production in B cells is tuned by mTORC1 signaling ^73,74^, and recent studies identify IgG accumulation as a key driver of ageing ^75,76^.

Lifespan extension in model organisms is generally associated with cost in reproduction ^52,77,78^. Notably, *Lsp2* mutant flies break this lifespan-reproduction tradeoff, which is unusual but not entirely impossible ^35,79,80^. This might arise from the fact that RPs are synthesized in great excess and probably never rate limiting for the assembly and functionality of ribosomes ^64^. In this sense, sparing the resources of translating excessive RPs should save unnecessary expenditure of cellular energy, yet without impacting normal animal physiology. This is further in line with a key notion in ageing that shifting cells from states of nutrient utilization to adult maintenance extends lifespan ^54^.

Ribosomes and translation have been center stage in lifespan regulation ^9,20,81^. The capacity and accuracy of protein synthesis declines with age, but manipulations that cause lifelong reductions in protein translation paradoxically extend lifespan in evolutionarily diverse organisms ^8,20,82,83^. Of note, evidence connecting translation and longevity initially came from yeast and *C. elegans* studies examining mutants of translation factors. However, the picture becomes more complex when it comes to higher eukaryotes, as genetic mutations in translational regulators and ribosomal components are often causally implicated in diseases ^84–86^, including ribosomopathies ^62^. Our serendipitous identification of a regulatory mechanism adjusting global RP translation urges us to search for similar regulators in mammals for lifespan extension.

How Lsp2 selectively affects RP translation remains an open question. We suspect that, besides the general translational repressor 4E-BP, this may also involve the RNA-binding protein LARP1, another mTORC1 substrate that directly binds TOP motif and controls RP production ^16,25,87,88^. Alternatively, Lsp2 may favor a specialized ribosome assembly that preferentially translates RPs, as ribosome heterogeneity does exist to fulfill specialized tasks of translation ^89,90^. Importantly, as the first *in vivo* demonstration of translational control of TOP genes, our study suggests *Drosophila* as probably the simplest model organism that has evolved functional TOP motifs in RP genes. Such an unanticipated similarity to mammals makes *Drosophila* a unique system to decipher the TOP biology from an evolutionary perspective and at the organismal level and reveal its implications for organismal metabolism and ageing.

## Methods

### Fly stocks and genetics

Fruit flies were cultured at 25℃ and under 60% humidity with a 12:12 light dark cycle. Conventional fly diet contains per liter 32.3g yeast, 69.2g corn flour, 9.2g soybean meal, 61.5mL syrup, 1.7g Nipagin methyl ester and 7.7g Agar. 3-5-day-old mated females were used for treatments unless otherwise noted. Experiments with mutant lines were performed at 25°C. For adult-onset expression, the TARGET system was used in combination with the indicated Gal4 drivers (*Lpp-Gal4*, *Mex-Gal4* and *Elav-Gal4*) to conditionally express UAS-linked transgenes ^91^. Flies were grown at 18-21°C to limit Gal4 activity. After 3 days at 18-21°C, adult progenies with the appropriate genotypes were shifted to 29°C, a temperature inactivating the temperature-sensitive Gal80’s ability to suppress Gal4. Gene-Switch system was also used for adult-specific gene manipulations ^92,93^, where 200 μM RU486 (Sigma #M8046) was supplemented to the fly medium and ingested by flies kept at 25°C. Key experimental conditions were indicated for other genetic crosses that used Gal4/UAS system for misexpression but did not involve *tub-Gal80^TS^*.

*Lsp2* stocks *UAS-Lsp2-GFP.attP40, UAS-Lsp2-3xHA.VK37, UAS-ΔSP-Lsp2-3xHA.VK37, Lsp2^B14^, Lsp2^A10^, Lsp2^KO^* were generated in this study (see next section for details). *Lsp2^G19255^* (BDSC27451) and P{Delta2-3}99B (P transposase line, BDSC3629) were used to generated *Lsp2^B14^*and *Lsp2^A10^* lines. *Lsp2^B14^* was used as genetic control of *Lsp2^A10^,* and both were crossed to *Thor^2^* to get *Thor*^2^; *Lsp2^B14^ and Thor*^2^; *Lsp2^A10^*lines*. Lsp2^KO^* was generated with CRISPR/Cas9 technique using *UAS-Cas9, nos-Gal4::VP16* (BDSC54593) as the source of Cas9 and backcrossed for seven generations into the *w^Dah^* background. *UAS-Lsp2; UAS-Lsp2* (Kyoto117651) obtained from Kyoto Stock Center was backcrossed for seven generations into the *w^1118^* (iso31) background. *Lsp2-Gal4* (BDSC6357) was used to visualize *Lsp2* expression by crossing to *UAS-mCD8::GFP* (BDSC32185).

4E-BP related fly stocks are *Thor-GFP^CPTI001137^* (Kyoto115070), *Thor^2^* (BDSC9559), *UAS-Thor-RNAi.KK* (VDRC100739) and VDRC-*KK* control line (VDRC60100). Drivers used are as follows. *tub-Gal80^TS^, UAS-GFP; Lpp-Gal4* (*Lpp^TS^,* from Bruno Lemaitre) was used as fat body specific driver. *tub-Gal80^TS^; Elav-Gal4* (*Elav^TS^, Elav-Gal4* from Liming Wang), *cg-Gal4* (from Lei Xue), *tub-GS^10^-Gal4* (from Fumiaki Obata) were used as additional drivers. A recombinant of *Lpp-Gal4, Lsp2^A10^* was made and further combined with *tub-Gal80^TS^*to get *tub-Gal80^TS^; Lpp-Gal4, Lsp2^A10^. Mex^TS^*(*Mex-Gal4, tub-Gal80^TS^*), *esg-Gal4* (from Bruno Lemaitre), *MHC-Gal4* (BDSC38464), *Elav-Gal4* (BDSC8765) and *Repo-Gal4* (from Christian Klaembt) on 2^nd^ chromosome were combined to *Lsp2^A10^.* The following fly stocks were also used in this study: *Oregon-R, w^1118^* (iso31) and *yw* used as wild type lines from Bruno Lemaitre; *w^Dah^* from Fumiaki Obata; *w^1118^* (iso CJ1) from Hongtao Qin; *UAS-InR^DN^* (BDSC8252), *UAS-PI3K92E-HA* (FlyORF001726), *UAS-PI3K^DN^* (BDSC25918), *UAS-PTEN* (BDSC82170), *UAS-AKT* (BDSC50758), *UAS-Rheb* (BDSC9688, isogenized to *w^1118^* (iso CJ1) line), *UAS-TSC1-RNAi* (VDRC22252), *UAS-RagA-B-RNAi* (TH201501134.S), *UAS-Rheb-RNAi* (BDSC33966), *UAS-mTOR-RNAi* (BDSC32941) to perturbate the Insulin and mTORC1 pathways; TRiP *attP2* control (BDSC36303), TRiP *attP40* control (BDSC36304), VDRC-*GD* control (VDRC60000) as RNAi control lines used to cross with Gal4 drivers; and *tub-Gal80^TS^* (BDSC7018 and BDSC7019).

### Generation of Lsp2 mutants, UAS-Lsp2-GFP, UAS-Lsp2-3xHA and UAS-ΔSP-Lsp2-3xHA lines

*Lsp2^B14^* and *Lsp2^A10^* were generated by mobilization of a P element inserted into the *Lsp2* locus. Essentially, *P{EP}Lsp2^G19255^*line was crossed to the *P{Delta2-3}99B* transposase and 100 candidate lines were created from the same genetic crosses that shared all the chromosomes. PCR screening was then performed to detect the absence of the original EP element and the *Lsp2* locus was sequenced for lines that had mobilized the P element. *Lsp2^B14^*was recovered as a precise excision therefore a revertant line used as our genetic control. *Lsp2^A10^* had 310 bp P element sequences left that contain a stop codon leading to premature stop of Lsp2 translation, thus an insertional mutant. We did recover several deletions with this method, but these deletions all extended to the neighboring gene *DEF8* therefore not being further used.

*Lsp2^KO^* flies were generated with the CRISPR/Cas9 method with two guide RNAs under the control of UAS ^47^ to target the beginning and the end of *Lsp2* coding sequence (CDS). The two gRNAs were synthesized and cloned into *pCFD6* vector using ClonExpress Ultra One Step Cloning Kit (Vazyme #C115). The resultant plasmid was verified by sequencing and further inserted into the *attP16* landing site by phiC31 integrase-mediated site-specific integration at UniHuaii (Zhuhai, China), to get the *UAS-Lsp2-sgRNA^2x^* transgenic line. This sgRNA line was crossed with a line stably expressing Cas9 in germline (BDSC54593), and *Lsp2^KO^* was obtained following screening with PCR. *Lsp2^KO^* was subsequently sequenced and found to have a deletion of 1,797 bp region (3L:12129587..12131384 in *D. melanogaster* (r6.62)) in *Lsp2* gene.

To generate the *UAS-Lsp2-GFP* fly line, the full-length *Lsp2* and *GFP* coding sequences were PCR amplified from *w^1118^*and a line carrying GFP, and cloned into *pUAST-attB* vector ^94^ using recombinant cloning method according to the ClonExpress Ultra One Step Cloning Kit. *GFP* was placed in frame with and downstream of *Lsp2* CDS. The resultant plasmid *pUAST-attB*-*Lsp2-GFP* was verified by sequencing. Transgenic fly was stablished by phiC31 integrase-mediated site-specific integration into the *attP40* site. To generate *UAS-Lsp2-3xHA* and *UAS-ΔSP-Lsp2-3xHA* lines, full-lengthen *Lsp2* CDS and *Lsp2* CDS lacking the 2^nd^-21^st^ codons (coding for its signal peptide) were assembled into the *pUAST-attB* vector together with the 3xHA sequences as a tag using the same method as for the generation of *UAS-Lsp2-GFP*. Both *UAS-Lsp2-3xHA* and *UAS-ΔSP-Lsp2-3xHA* were inserted into the *VK37* site by UniHuaii (Zhuhai, China). Primers used for cloning are available upon request.

### Generation of Lsp2 antibody

Lsp2 antibody was generated by immunizing rabbits with a purified Lsp2 fragment by Mabnus Biotech (Wuhan, China). C-terminal protein sequences 461-701 of Lsp2 was expressed by HEK 293F and purified by Nickel column. Purified Lsp2 protein was used to immunize rabbits via 4 injections, with the interval of 28 days, 14 days, and 14 days between injections. Freund’s complete adjuvant (FCA) was used in the first injection, and Freund’s incomplete adjuvant (FIA) for the rest injections. 3 days after the 4^th^ injection, antiserum titer was tested by ELISA. The rabbits with higher titer were killed and bled at the 64^th^ day since the first injection. The affinity column was made by coupling 1mg purified Lsp2 protein to CNBr-activated Sepharose 4B from GE Healthcare. The antiserum was applied and bound to the column, and then the specific Lsp2 antibody was eluted with the Glycine HCl buffer of pH 2.5, followed by dialysis and concentration.

### Dietary manipulation and drug treatments

For yeast extract assays, flies were fed with 1% sucrose and different concentrations of yeast extract from Oxoid™ Yeast Extract Powder (Thermo Scientific #LP0021B) for 48h. The following amino acids were purchased from Sigma and used to induce *Lsp2* expression: L-alanine (#A7627), L-arginine (#A5131), L-asparagine (#A0884), L-aspartate (#A9256), L-cysteine (#C1276), L-glutamate (#G1251), L-glutamine (#G3126), L-glycine (#G7126), L-histidine (#H8000), L-isoleucine (#I2752), L-leucine (#L8000), L-methionine (#M9625), L-phenylalanine (#P2126), L-proline (#P0380), L-serine (#S4500), L-threonine (#T8625), L-tryptophan (#T0254), L-tyrosine (#T3754), L-valine (#V0500), and L-lysine (#L5626). They were soaked to a filter paper and used to feed flies for 24-48 h at 5% w/v diluted in water containing 1% sucrose. When applying multiple AAs simultaneously, we took equal amount for each AA and the total amount of all AAs used summed to 5% w/v in water containing 1% sucrose. In addition, feeding experiments were also performed with amino acids at concentrations specified in the FLYAA recipe ^95^.

Sugar-Yeast-Agar (SYA) foods were used to analyze fly lifespan response to varying protein concentrations. SYA foods contained 5% sucrose, 1.5% agar and different levels of yeast (2.0X SYA containing 20% yeast; 1.0X SYA containing 10% yeast; 0.5X SYA containing 5% yeast; 0.1X SYA containing 1% yeast; and 0X SYA containing no yeast). SYA foods also contained the same concentration of Nipagin methyl ester as in our conventional fly medium. Yeast used was golden version of instant dry yeast from Angel Yeast (Wuhan, China). Except for lifespan experiments, flies were allowed to ingest SYA foods for 7 days before other tests were performed.

For rapamycin assays, 200 μM rapamycin (LC Laboratories #R-5000) was supplemented to the foods as indicated in Figures. For puromycin incorporation assays, females were transferred into 1.0X SYA diet supplemented with 600 μM puromycin (AbMole #M3637) for 24h before further analyzed for global protein synthesis.

### Lifespan

Flies were reared at a standard density prior to lifespan experiments. Genetic crosses were performed at the indicated temperature, and the progenies were collected and allowed to mate for 3 days. Flies were then maintained in vials containing respective culturing medium at a density of 20 flies per vial. Flies were transferred to fresh food vials every two days and the number of deaths was scored daily. Survival curves were generated and analyzed using GraphPad Prism 9 software, and representative results were shown in Figures.

### Food intake

Without performing prior starvation that interrupts *Lsp2* expression and normal physiological states of the animal, flies sorted 3-4 days in advance under CO_2_ anesthetics were transferred to new food vials (20 flies/vial) containing 0.5% FD&C Blue No. 1 (Sigma #861146) in conventional fly diet for 24 h. Note that this blue dye is not absorptive. Flies were then flash frozen in liquid nitrogen for 30 s and the heads were removed by vigorous shaking. The remaining parts of the flies were collected in tubes with 600 μL of PBS and homogenized with the Precellys Evolution Homogenizer (Bertin technologies), followed by centrifugation at 15,900 × g for 25 minutes (min). 100 μL supernatants was added to a 96-well plate and the absorbance was measured at 620 nm with the Synergy HTX (BioTek) multimode plate reader. Absorbance was calculated to represent levels of food intake. At least 100 flies were used for each genotype. To infer food intake by measuring excreta ^96^, flies were transferred into a food vial (5 flies/vial) containing 0.5% blue dye (Sigma #861146) for 24 h followed by recording the number of deposits.

### Developmental timing and fecundity

Adult females were fed fresh yeast for 2 days and allowed to lay eggs for 4h on apple juice plates (1/3 apple juice and 2/3 water with 2% agar) with yeast paste. About 40 first instar larvae were collected and placed into new food vials 26 h later. Then the developmental timing was monitored at 25°C by counting the number of pupariation and eclosion every 2-3 h. For fecundity assays, adult females were fed fresh yeast for 24 h after a dry starvation in an empty vial for 6 h. Then, the flies were flipped into new food vials (5 females per vial) containing a basic diet (1% yeast, 5% sucrose and 1% agar) and monitored every 24 h for egg laying for 5 days. Lifelong fecundity test was done as follows. 3 females and 3 males that had just eclosed 1 day before were sorted into conventional food vials and monitored for the number eggs laid every two days for 30 days; alternatively, 5 newly eclosed females and males were put together and the number of eggs laid were counted after 10 and 30 days.

### Starvation, oxidative stress, and systemic bacterial infection

For starvation assays, 5-day-old female or male flies were put on 1% agar tube essentially with only the access to water, and their survival was recorded. For oxidative stress assays, 5-day-old female flies were fed the 1.0X SYA diet supplemented with 17.5 μM paraquat (Sigma #856177), and their survival was monitored over time. Systemic infection was performed by pricking the thorax of 10-or 30-day-old adult females with a 100 μm-thick insect pin dipped into a concentrated pellet of *Providencia rettgeri* (*P. rett*, from Bruno Lemaitre) at an optical density (OD_600_) of 5, and survival was recorded post-infection at 29°C.

### Locomotor activity

Climbing. Flies were collected and further reared to the indicated age before being assayed for climbing ability (negative geotaxis). Female flies were anesthetized with CO_2_ two days prior to the experiment and divided into groups of 10 flies/tube. For the test, the flies were placed in an empty fly vial (height: 9.5cm; diameter: 2.4cm) marked with lengthen scales in 1cm increment and allowed to acclimate for 30 min. Then, we gently tapped the flies to the bottom of the tube and immediately initiated video recording of fly climbing. This was repeated three times with a 5-min interval between trials. From the video recordings, the distance of each fly climbed for 5 s (for 10-day-old females) or 10 s (for 30-day-old females) was determined, and the average climbing distance of 10 flies in the same vial was calculated and was further averaged from the three technical repeats. At least three biological replicates were conducted for each experiment. The climbing assays were performed at room temperature. (2) Spontaneous walking. Video tracking was used to assess spontaneous locomotion. Flies were anesthetized on ice and loaded 1 fly/well into a 48-well plate. The bottom of the 48-well plate was filled with 1% agar and 5% sucrose so that flies can only move horizontally. After a 2 h recovery period at room temperature, the flies were placed into the Zebrabox tracking system (ViewPoint) to track fly movement. From the tracking data, we calculated the velocity and total distance travelled by each fly over an 8-hour period. The tests were routinely done from ZT3-11 (10am-6pm) at room temperature, and the test plates were illuminated during tracking. All experiments were performed in biological triplicate.

### qPCR

Total RNA was extracted from head or body of 15-20 flies using RNAiso Plus (TaKaRa #9109). RNA concentration was measured by Nanodrop One (Thermo Fisher Scientific). cDNA was synthesized with oligo dT using the PrimeScript™ RT Reagent Kit (TaKaRa #RR037A) using 200 ng of total RNA as template, followed by 20-fold dilution of the resulting first strand cDNA with nuclease-free water prior to quantitative real-time PCR analysis. qPCR was performed in at lease duplicate for each sample using SYBR Green (Roche #04887352001) on a q225 qPCR System from Quantagene (Kubo Technology). Expression values were calculated using the ΔΔCt method and relative expression was normalized to *RpL32*. The expression in control sample was further normalized to 1. For qPCR results, each circle shown in figures represents one independent biological replicate. Of note, as *Lsp2* coding sequences almost completely overlap with a non-coding RNA (*CR11538*), we designed primers that amplify the 3′ UTR of *Lsp2* transcripts, regions specific to *Lsp2*. Primer sequences are as follows.

*Lsp2-3′UTR*: 5′-*CGGTGAAGCATGACTACTACTT*-3′ and 5′-*TTCGTTGGGTCCAGGATCTAG*-3′; *Lsp2*: 5′-*ATGAAGTCGTTCACGGTGAT*-3′ and 5′-*CTAGACCACATTGGTGTGCT*-3′; *RpL32*: 5′-*TCTGCATGAGCAGGACCTC*-3′ and 5′-*ATCGGTTACGGATCGAACAA*-3′; *Lsp1α*: 5′-*GCGGGATCCCATGTTCTATATG*-3′ and 5′-*GAAGTTGCTCCTTGGTGTATTTG*-3′; *Lsp1β*: 5′-*GACCTACGCCAACTACGATATG*-3′ and 5′-*CGGTCTTGTAGTAGCCCATTT-*3′; *Lsp1γ*: 5′-*ATCCGACTTCTTTGTG CCTAAT*-3′ and 5′-*CGAACTGGGTGTACTTCTTCTC*-3′; *dilp2*: 5′-*AGCAAGCCTTTGTCCTTCATCTC*-3′ and 5′-*ACACCATACTCAGCACCTCGTTG*-3′; *Hsp70Ba*: 5′-*GTTGAAAGTATTCTCTTCTTG*-3′ and 5′-*CCAAGTAAATCAACTGCAAC*-3′; *Thor*: 5′-*AGTTCTTAGATGCACCCACTAAA*-3′ and 5′-*GGGTCAATATGACCGAGAGAAC*-3′.

### Immunoblotting

Tissues were homogenized and lysed in the 0.5% Triton X-100 buffer with the addition of 2× cOmplet Protease Inhibitor Cocktail (Roche #11697498001) and Phosphatase Inhibitor Cocktail I and Ⅱ (MCE #HY-K0021 and #HY-K0022) at 4℃. Protein samples were analyzed by SDS-PAGE and transferred to 0.22μm polyvinylidene difluoride (PVDF) membranes, using standard Western blotting procedures. Total protein loading was visualized via Ponceau S (Coolaber #SL1281) staining for puromycin incorporation assays. Membranes were blocked in 5% non-fat milk (Coolaber #CN7861) for at least 60 min at room temperature and probed with primary antibodies overnight at 4°C. The following primary and secondary antibodies were used: rabbit anti-Lsp2 (this study; 1:2,000), rabbit anti-p-S6K (CST #9209; 1:1,000), rabbit anti-p-RpS6 (produced in Yan Yan lab; 1:10,000), rabbit anti-p-eIF2α (CST #3398; 1:1,000), rabbit anti-p62 (gift from Hongtao Qin; 1:50), rabbit anti-p-4E-BP (Thr37/46) (CST #2855; 1:500), rabbit anti-non-phospho-4E-BP1 (Thr46) (CST #4923; 1:500), mouse anti-RpS6 (CST #2317; 1:500), rabbit anti-RpS15A (ABclonal #A10241; 1:1,000), rabbit anti-RpL30 (ABclonal #A13690; 1:1,000), rabbit anti-RpL40 (Abcam #ab109227; 1:1,000), mouse anti-GFP (Proteintech; 1:10,000); rabbit anti-HA (CST #3724; 1:1,000), mouse anti-Actin (Bioss #bsm-8777M; 1:1,000), mouse anti-Tubulin (DSHB #4A1; 1:500), mouse anti-Puromycin (Sigma #ZMS1016; 1:1,000), goat anti-rabbit IgG (Abbkine #A21020; 1:8,000) and goat anti-mouse IgG (Abbkine #A21010; 1:8,000). Detection was performed using ChemiDoc MP Imaging system (Bio-Rad). Densitometric analysis of blot images was carried out using ImageJ (version 1.52a) and the gray scale of the control sample was normalized to 1.

### Immunofluorescence

Flies were dissected in PBS and fixed for 60 min with 4% paraformaldehyde (EMS #157-8) in PBS. Samples were rinsed in PBS with 0.1% TritonX-100 (PBT) and blocked with 2% BSA in PBT for at least 60 min at room temperature, followed by primary antibody incubation in 2% BSA PBT at 4℃ overnight. Then samples were washed 3 times in PBT and incubated with secondary antibodies and DAPI (Sigma #D9542; 1:10,000) for 2 h at room temperature. Samples were washed 3 times in PBT, mounted and imaged using Zeiss Imager M2 microscope. Quantification of Dilp2 staining intensities in the brain IPCs was performed using ZEN software (version 2.3). The following primary and secondary antibodies were used: rabbit anti-Dilp2 (gift from Jan Veenstra; 1:1,000), rabbit anti-Lsp2 (this study; 1:100), Chicken anti-GFP (Abcam #ab13970; 1:1,000), rabbit anti-Phospho-Histone H3 (CST #9701; 1:500), Goat anti-Chicken IgY (H+L)Alexa Fluor™ 488 (Invitrogen #A11039; 1:1,000), Goat anti-Rabbit IgG (H+L) Alexa Fluor™ 488 (Invitrogen #A11008; 1:1,000), Goat anti-Rabbit IgG (H+L) Alexa Fluor™ 568 (Invitrogen #A11036; 1:1,000).

### ELISA

Haemolymph was extracted from 50 decapitated females/sample by centrifugation 1.5mL tube with Nylon mesh filter at 1,500 × g for 15 min at 4°C. Then, 0.5 μL haemolymph was diluted in 50 μL PBS and added into ELISA plate for overnight at 4°C. After being washed 4 times with PBS, the samples were blocked in 5% BSA at 4°C for overnight, followed by incubation with primary antibody anti-Dilp2 (gift from Jan Veenstra, 1:2,000) in 1% BSA for 2 h at room temperature. Then samples were incubated with the secondary antibody Goat anti-Rabbit IgG HRP (Abbkine #A21020; 1:5000) for 2 h at room temperature after being washed 5 times with 0.2% PBT. The ELISA plate was further washed 10 times with 0.2% PBT and added 200 μL TMB (Sino Biological #SEKCR01). After a 10-min incubation, 100 μL Stop Solution (Solarbio #C1058) was added to the ELISA plate and the absorbance was measured at 450 nm with the Synergy HTX (BioTek) multimode plate reader.

### RNA sequencing and data analysis

This study involves three regulator transcriptomic analyses (RNA-seq) using the heads of flies with the following detailed specifications. (i) Diet group: 7-days-old *w^1118^*(iso31) female flies raised on conventional lab diet and further fed for 2 days with either a control diet (2% sucrose) or a protein-rich diet (2% sucrose + 3.3% yeast extract) at 25°C; (ii) Age group: 3-day-old (as Young) and 50-day-old (as Old) *Oregon-R* flies raised on conventional lab diet at 25°C; (iii) *Lsp2* and *Rheb* overexpression: flies with fat body specific overexpression of *Lsp2* or *Rheb* by *Lpp^TS^* and respective control flies raised on conventional lab diet for 3 days and then fed with a sugar-only diet (5% sucrose) for 2 days at 29°C.

Total RNA was extracted from heads of females for RNA-seq analysis. Fly heads were dissected in ice-cold PBS, transferred into RNAiso Plus reagent (TaKaRa #9109), and were homogenized with the Precellys® Evolution Homogenizer (Bertin technologies). 20 heads were used for each sample in biological triplicates. Total RNA extraction was performed as previously described ^97^. RNA concentration was measured by Nanodrop One (Thermo Fisher Scientific). RNA integrity was assessed using the Agilent 2100 Bioanalyzer (Agilent Technologies). Then, the sequencing libraries were constructed using VAHTS Universal V10 RNA-seq Library Prep Kit (Vazyme) and sequenced on an Illumina Novaseq 6000 platform with 150 bp paired-end reads. Raw reads of FASTQ format were firstly processed using fastp and the low-quality reads were removed to obtain the clean reads that were mapped to the reference genome using HISAT2. FPKM of each gene was calculated and the read counts of each gene were obtained by HTSeq-count. Principal component analysis (PCA) analysis was performed using R (v3.2.0). Differential gene expression analysis was performed using the DESeq2. We set q value < 0.05 and foldchange > 2 or foldchange < 0.5 as the threshold for significantly differential expression gene (DEGs). Unsupervised hierarchical clustering of DEGs was implemented in R (v3.2.0) to visualize expression patterns across samples via heatmaps. Functional enrichment analyses of DEGs, including Gene Ontology (GO) and Kyoto Encyclopedia of Genes and Genomes (KEGG) analyses, were performed to identify significantly enriched biological terms or pathways. The transcriptome sequencing and analysis were conducted by OE Biotech (Shanghai).

### Proteomics

Five biological replicates were included for both *Lsp2^B14^* and *Lsp2^A10^* flies. 80 flies/sample were fed the 0.5X SYA diet for 14 days, collected and ground into powder using liquid nitrogen. Approximately 20 mg of powder was resuspended in 200 μL lysis buffer (4% SDS, 100 mM DTT, 150 mM Tris-HCl pH 8.0). The samples were boiled and further ultrasonicated. Undissolved cell debris were removed by centrifugation at 16,000 x g for 15 min. The supernatant was collected and quantified with a BCA Protein Assay Kit (Bio-Rad). Protein digestion was performed using the FASP (Filter-Aided Sample Preparation) procedure ^98^. The digested proteins were transferred into 1.5-mL Nanosep tubes and centrifuged for 20 min at 14,000 × g at 20 °C to remove the DTT. Then, 100 μL 50 mM iodoacetamide (IAA) in UA buffer (8M urea and Tris-HCl) was added to block reduced cysteine residues through a 20-min-incubation in darkness. Then, 200 µL ammonium bicarbonate (100mM) buffer was added to the beads in the filter cartridge, centrifuged for 15 min at 14,000 × g at 25°C. Finally, 50-100 μL of 50 mM ammonium bicarbonate buffer with trypsin was added at the enzyme: protein ratio of 1:50 into the sample and incubated at 37℃ for 20 h. The peptides were harvested by centrifugation, acidified by 1% formic acid (FA) and subsequently dried by a refrigerated CentriVap Benchtop Vacuum Concentrators (Labconco, Kansas, MO). Finally, the trypsin-digested peptides were desalted with C18 StageTip. The concentrations of peptides were determined with OD_280_ by Nanodrop One.

LC-MS were performed on a Q Exactive Plus mass spectrometer coupled with an EASY-nLC 1200 nano-liquid chromatography (nLC) system (Thermo Fisher Scientific) by Shanghai Bioprofile Biotechnology, according to the standard MS procedure. MS data was acquired using a data-dependent top20 method that dynamically chose the most abundant precursor ions from the survey scan (300–1800 m/z) for higher-energy collisional dissociation (HCD) fragmentation. The instrument was run with the peptide recognition mode enabled. A lock mass of 445.120025 Da was used as internal standard for mass calibration. The full MS scans were acquired at a resolution of 70,000 at m/z 200, and 17,500 at m/z 200 for MS/MS scan. The maximum injection time was set to for 50 ms for MS and 50 ms for MS/MS. Normalized collision energy was 27 and the isolation window was set to 1.6 Th. Dynamic exclusion duration was 60 s. The MS data were analyzed for protein identification against the *Drosophila melanogaster* protein database at https://www.uniprot.org/taxonomy/7227. The database search results were filtered and exported with <1% false discovery rate (FDR) at peptide-spectrum-matched level, and protein level, respectively. Label-free quantification was carried out in MaxQuant using intensity determination and normalization algorithm as previously described ^99–101^. The “LFQ intensity” of each protein in different samples was calculated as the best estimate, satisfying all of the pairwise peptide comparisons, and this LFQ intensity was almost on the same scale of the summing peptide intensities. The quantitative protein ratios were weighted and normalized by the median ratio in MaxQuant software. Only proteins with fold change ≥1.5-fold and p value <0.05 were considered as significantly differentially expressed proteins.

### Polysome profiling and deep sequencing of fractioned mRNAs

Polysome profiling was performed as described with a slight modification ^102^. 15-day-old female flies (∼60/sample) were anesthetized on ice and the whole flies were ground in liquid nitrogen. A portion of the powdered tissue was set aside for parallel mRNA-seq, while the remaining was resuspended in lysis buffer (50 mM Tris-Cl, pH 7.5, 150 mM NaCl, 5 mM MgCl₂, 1% Triton-X 100, 2 mM DTT, 500 U/mL Superase In RNase Inhibitor, 20 µg/mL Emetine, and 50 µM GMP-PNP), supplemented with one tablet of EDTA-free cOmplete Protease Inhibitors (Roche #4693132001) per 5mL. The samples were thoroughly homogenized using a homogenizer to ensure complete tissue lysis. Then, the samples were cleared by centrifugation at 14,000 × g for 5 min at 4°C. The supernatant was quantified using a Qubit 4 fluorometer (Thermo Fisher Scientific). Aliquots of 50 µl of the supernatants were used for the construction of parallel mRNA-Seq libraries. Equal amounts of lysate from different samples were then loaded onto a 10%-45% sucrose density gradient (50 mM Tris-Cl, pH 7.5, 250 mM NaCl, 15 mM MgCl₂, 0.5 mM DTT, 40 U/µL RNase OUT, and 20 µg/mL Emetine) in a SW41Ti rotor, supplemented with half a tablet of EDTA-free cOmplete Protease Inhibitors (Roche #4693132001) per 50 mL, and centrifuged at 35,000 rpm for 3 h at 4°C. After ultracentrifugation, the gradients were fractionated using the Gradient Profiler (BioComp Instruments), with continuous monitoring of absorbance at 260 nm. The monosome and polysome fractions were collected for subsequent construction of translatome libraries. Polysome profiling was performed at Analytical Instrumentation Center of Hunan University.

RNA from the monosome and polysome fractions was separately collected using TRIzol reagent (Invitrogen #15596026) for subsequent construction of translatome libraries. RNA from all three sources (total RNA, monosome-bound RNA and polysome-bound RNA) was processed identically as follows. Poly(A)^+^ RNA was enriched from 2 µg of total RNA or translating RNA using Library Preparation VAHTS mRNA Capture Beads (Vazyme #N401), followed by RNA fragmentation, end repair, adaptor ligation, and PCR amplification with the VAHTS Universal V6 RNA-seq Library Prep Kit for Illumina (Vazyme #NR604). After quality control and quantification, the prepared libraries were sequenced on Illumina NovaSeq 6000 platform, generating paired-end reads of 150 bp.

Sequencing data were processed as described previously ^103^. All reads were aligned to the *Drosophila melanogaster* (BDGP6.46) reference genome by STAR. Uniquely mapped reads were summarized to *Drosophila_melanogaster*.BDGP6.46.110.chr.gtf annotated genes using feature Counts. The resulting raw gene expression matrix was then imported into the R environment for further processing and analysis. Differential expression analyses were performed using R package DESeq2 based on an adjusted p value < 0.05 and absolute value of log_2_ (Fold Change) > 0.5. For transcriptome analysis, differentially expressed genes (DEGs) were identified using mRNA-seq data. Translation efficiency (TE) for differentially translated transcripts was further determined based on monosome-seq, polysome-seq and mRNA-seq data. Using polysome-seq data as the translatome data and mRNA-seq data as the transcriptome data, TE of each gene was calculated through the deltaTE method ^104^. Gene ontology analysis of significantly regulated transcripts was done with Cluster Profiler (4.10.1).

### TOP mRNA identification and TOP motif identification

The TOP score for each protein-coding transcript annotated in FlyBase r6.06 was calculated based on the nucleotide composition within the first eight nucleotides, following a previous described method ^102^. A total of 112 5′TOP genes were identified in *Drosophila* (TOP score >3), and were collectively referred to as fly TOP genes. Ribosomal protein-coding genes for each species were retrieved from the KEGG database in JSON format, and their cDNA sequences were downloaded from the NCBI database. The first 20 nucleotides of each RP’s cDNA were extracted to generate sequence logos using the ggseqlogo (v0.2) for visualization ^105^. To characterize regulatory motifs, the 5’ end sequences of transcripts were examined, and the following elements were annotated (1) the TOP motif, defined as a cytosine followed by 4-20 consecutive pyrimidines at the extreme 5’ end; (2) the *iso-TOP*: defined as a Uracil followed by 4-20 consecutive pyrimidines at the extreme 5’ end; and (3) the pyrimidine-rich sequences, defined as a region near the 5′ end containing at least six consecutive pyrimidines while allowing no more than one purine. These motifs were manually annotated in mRNAs translationally downregulated following *Lsp2* loss, according to the abovementioned criteria.

### Structure modeling

The protein sequences of Thor (UniProt ID: Q9XZ56) and Lsp2 (UniProt ID: Q24388) from *Drosophila melanogaster* were obtained from the UniProt (https://www.uniprot.org/). The predicted models of Thor and Lsp2 were generated using the AlphaFold 3 online platform (https://alphafoldserver.com/) by employing the pairformer and diffusion modules. The Lsp2 monomer-Thor complex and Lsp2 hexamer-Thor complex were generated using the built-in interaction prediction module of AlphaFold 3 with default parameters. The highest-confidence model was selected based on the pLDDT score. Structures were visualized with PyMOL (https://pymol.org/2/) to further analyze the protein-protein interaction interface. Key residues at the interaction interface were highlighted to assess their potential role in the stability and function of the protein complex.

### Statistical analyses

Quantitative data are shown essentially as mean ± s.e.m. from at least three biological replicates of each experiment. For all quantifications, n represents the number of biological replicate. In Figures S2q-r (S stands for “Extended Data Fig.”), S3c-d (Speed), S9k and S9n, one dot represents one fly. In Figures S3c-d (*middle left*), one dot represents the average distance climbed by 10 flies in 5 or 10 seconds. In Figures 1c-f, 1h-i, 1k-l, 1o, 4c, 4f, S1c-d, S1h-m, S2b, S4f-g, S9c, and S9i, one dot represents 20 heads. Other images shown in Figures to characterize gene expression/protein localization are representative of at least 20 individuals and the experiments were repeated at least twice. Statistical significance was determined using either the unpaired *t* test for comparison of two groups, or one-way ANOVA with Tukey’s multiple comparisons test or Dunnett’s multiple comparisons test where multiple comparisons were necessary, in GraphPad Prism Software (version 9.5.1). Survival data were pooled and analyzed in Prism software using the Log-rank test (Figures 2b, 2d, 4a, 4h, 4j, 5a-c, S2f-p, S3a-d, S5g and S5l). *P* values are indicated in Figures. Complete details regarding the statistical tests used, sample sizes, and levels of significance are reported in the Figures and figure legends.

## Data availability

All relevant data are available from the corresponding authors upon reasonable request. Regular RNA-seq data are deposited in GEO under the accession numbers GSE295794 (age group; diet group; *Lsp2* and *Rheb* overexpression). The mRNA-seq, monosome-seq and polysome-seq data related to polysome profiling are available under the accession number GSE297586. The proteomic data have been deposited to the ProteomeXchange Consortium via the iProX partner repository with the identifier PXD063648 (IPX0011855000).

## Acknowledgements

We would like to thank Drs. Claudine Neyen, Sebastian Sorge and Fumiaki Obata for comments on the manuscript; Drs. Jian Qiu, Zheng Guo, and Dingxiao Zhang for valuable suggestions; Drs. Bruno Lemaitre, Youheng Wei, Jan Veenstra, Jiong Chen, Hongtao Qin and Yan Li for kindly sharing fly strains, antibodies, and bacterial strains; BDSC, VDRC and Kyoto Stock Center for fly stocks; DSHB for antibodies; and Zhai Lab members Jianhan Yi, Jingyun Su and Xinfeng Wang for technical support. This work was supported by Hunan NSF grants 2025JJ30009 (to Z.Z.) and 2025JJ40028 (to Y. Wang); NSFC grants 32170509 (to Z.Z.), 32470655 (to Y.Wang) and 32371219 (to J.L.); the Key Project of Developmental Biology and Breeding from Hunan Province 2022XKQ0203 (to Z.Z.), Shanghai Pujiang Program 23PJ1415500 (to J.L.), and Hong Kong Grant Council GRF16103620 and GRF16104324 (to Y.Y.).

## Author contributions

J.W. and Z.Z. developed the research plan and together with J.L. and Y.Wang interpreted experimental results. J.W. designed and performed all the fly work. Z.C. performed polysome profiling, associated RNA sequencing and TOP motif analysis. J.G. and K.C. helped with qPCR, lifespan, infection, and food intake analyses. M.Y. helped with analysis of regular deep sequencing data. X.N. helped with polysome profiling. Y.Wen helped with protein structural modeling. Y.Y. provided a key reagent. Y.Wang and Z.Z. co-supervised the project. Z.Z. wrote the manuscript. All authors read and approved the final manuscript.

## Competing interests

The authors declare no competing interests.

Correspondence and requests for materials should be addressed Zongzhao Zhai.

---

**Extended Data Fig. 1.**
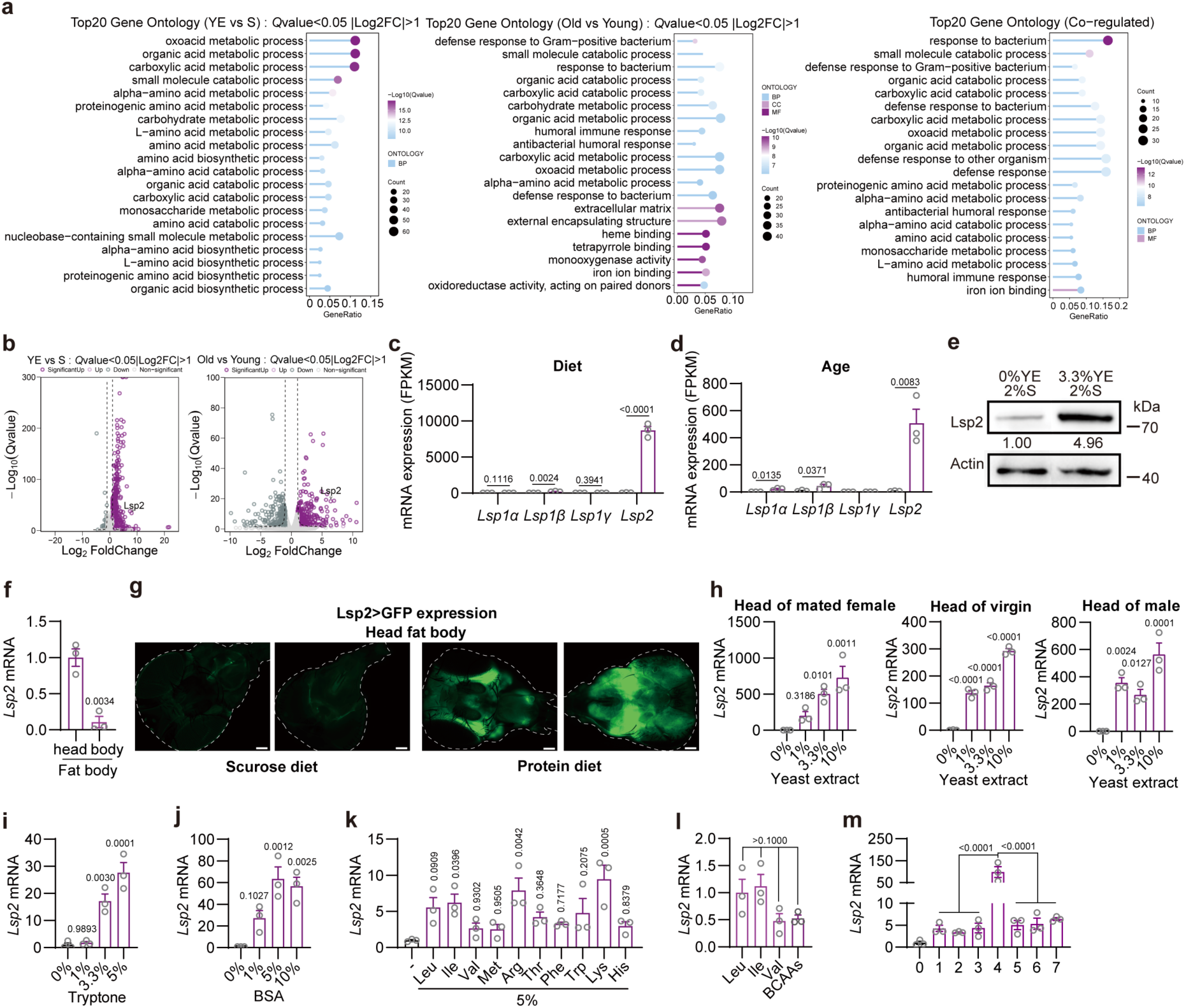
*Lsp2* is induced by balanced dietary EAAs. **a**, Gene ontology (GO) analyses of significantly differentially expressed genes (DEGs, *Q*<0.05 & |log_2_FoldChange|>1) in the YE vs S comparison (Diet), the Old vs Young comparison (Age) and the 309 co-regulated DEGs in both datasets. Shown are top 20 significantly enriched GO terms. **b**, Volcano plots of upregulated DEGs in both datasets with *Lsp2* indicated. **c**,**d**, Expression of the four *LSP* genes (*Lsp1α*, *Lsp1β*, *Lsp1γ* and *Lsp2*) in the transcriptomic datasets. **e**, Western blots showing Lsp2 protein levels in whole flies of 5-day-old *w^1118^* females treated with the indicated diets for 2 days. **f**, Relative *Lsp2* mRNA levels in dissected fat body from the heads and from the bodies of 7-day-old *w^1118^*females raised on our conventional lab diet. *Lsp2* expression in the head fat body is normalized to 1. **g**, *Lsp2* expression in fly heads indicated by *Lsp2-Gal4>UAS-GFP*. 5-day-old females were fed 1% sucrose or 1% sucrose with 10% yeast extract (YE) for 2 days. **h**, Relative *Lsp2* mRNA levels in heads of 5-day-old mated *w^1118^*females, virgin females, and males raised on the indicated diets for 2 days. All diets contained 1% sucrose. *Lsp2* expression on sucrose-only food is normalized to 1. **i**,**j**, Relative *Lsp2* mRNA levels in heads of 5-day-old mated *w^1118^* females raised on the indicated diets of alternative protein sources for 1 (**j**) or 2 (**i**) days. 1% sucrose was added in **i** and 5% sucrose in **j**. *Lsp2* expression on sucrose-only food is normalized to 1. **k**-**m**, Relative *Lsp2* mRNA levels in heads of 5-day-old mated *w^1118^*females raised on the indicated amino acid (AA) diets with 1% sucrose for 1 day. *Lsp2* expression on sucrose-only food is normalized to 1 in **k**,**m**. The AA contents sum to 5% w/v in **k**-**m**, and each AA shares equal weight ratio when multiple AAs are included in one diet (**l**,**m**). BCAA is a mixture of Leu, Ile and Val. In **m**, the numbers under the *x* axis stand for Leu, Ile and Val (**1**, BCAAs); Leu, Ile, Val, Phe and Trp (**2**, BCAAs + Phe + Trp); Leu, Ile, Val, Phe, Trp, Lys and Arg (**3**, 7 EAAs); Leu, Ile, Val, Phe, Trp, Lys, Arg, Met, Thr and His (**4**, a full EAA diet); Phe, Trp, Lys, Arg, Met, Thr and His (**5**, full EAAs lacking BCAAs); Lys, Arg, Met, Thr and His (**6**, 5 EAAs); Met, Thr and His (**7**, 3 EAAs). 20 females were used for each sample and represented as one dot in **c**,**d**,**f**,**h**-**m**, and these data are mean ± s.e.m. Shown in **e,g** are representative images of three biological replicates. Student’s *t*-test in **c**,**d**,**f**; one-way ANOVA with Dunnett’s multiple comparisons test in **h-k,m**; one-way ANOVA with Tukey’s multiple comparisons test in **l**. *P* values are indicated. Scale bars 50 µm in **g**.

**Extended Data Fig. 2.**
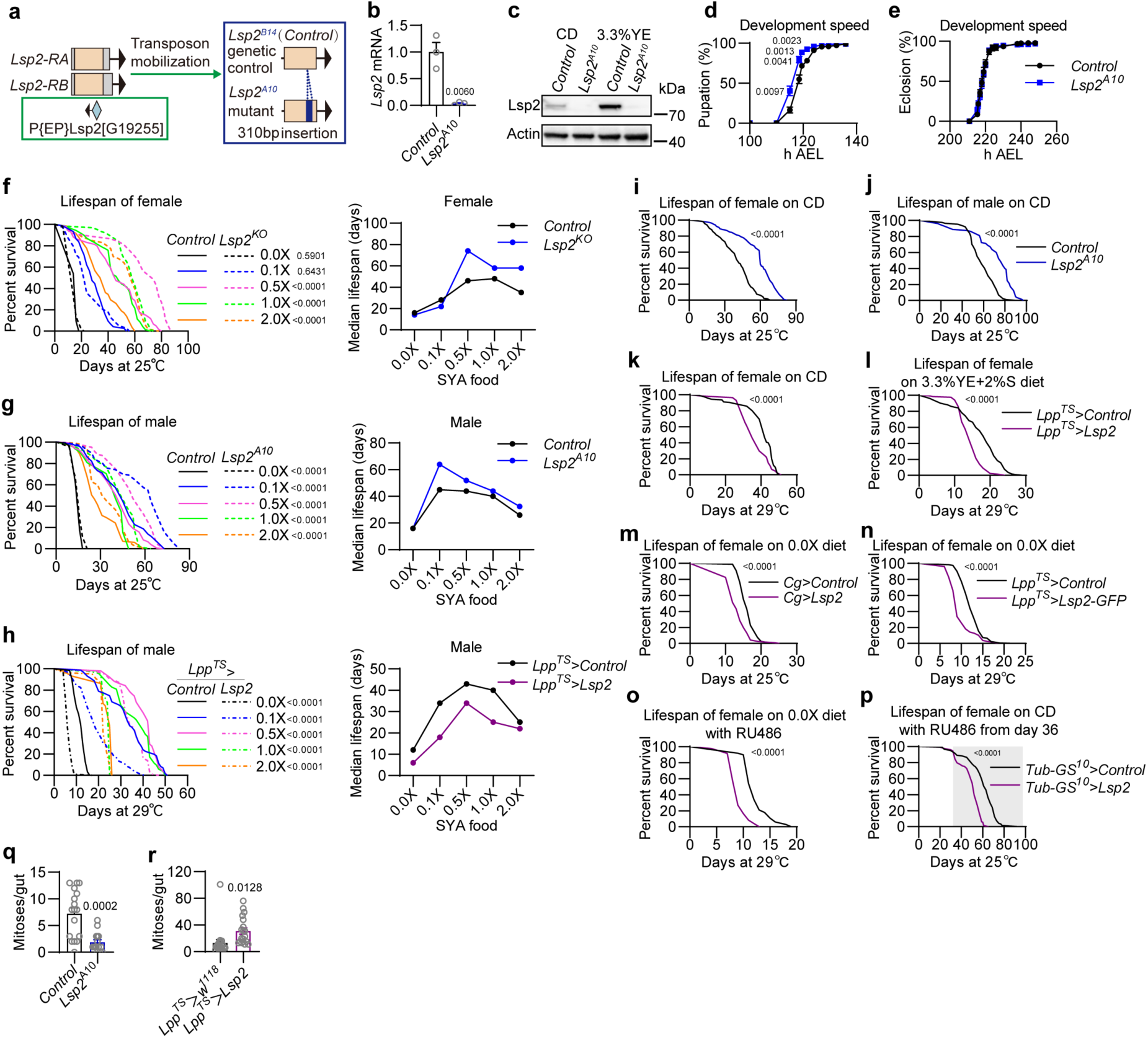
Lsp2 regulates lifespan. **a**-**c**, Schematic (**a**) of the generation of *Lsp2^A10^* mutant line and its genetic control *Lsp2^B14^*by mobilizing the indicated *P{EP}* transposon, and validation of the lines by qPCR (**b**) and Western blots (**c**) using the heads (**b**) or whole bodies (**c**) of 5-day-old females raised for 2 days on conventional diet (CD; **b**,**c**) and a diet consisting of 3.3% YE and 2% sucrose (**c**). **d**,**e**, Developmental timing analyses of *Lsp2^A10^* flies and its genetic control on CD. Pupation time (**d**) and adult eclosion time (**e**) are indicated as hours taken after egg laying (AEL) in a group of 40 synchronized 1^st^ instar larvae. n = 5 for *Control*; n = 6 for *Lsp2^A10^*. **f**-**h**, Survival (*left*) and median lifespan (*right*) of flies with the indicated genotypes and sexes on the five SYA diets. Numbers of flies used in 0.0X, 0.1X, 0.5X, 1.0X and 2.0X SYA foods, respectively, are 63, 74, 59, 70 and 58 for the *Control* of *Lsp2^KO^*; 57, 39, 37, 63 and 53 for *Lsp2^KO^*; 115, 101, 120, 78 and 99 for the males of *Lsp2^A10^ Control* line; 117, 120, 104, 94 and 106 for the males of *Lsp2^A10^*; 61, 71, 51, 52 and 43 for the males of *Lpp^TS^>Control* line; 94, 96, 93, 94 and 83 for the males of *Lpp^TS^>Lsp2* line. **i**-**p**, Survival analyses of female flies (except **j**) with the indicated genotypes raised on the indicated diets. 200μM RU486 was added either throughout the analysis (**o**) or from the middle life at day 36 onwards (**p**) to activate transgene expression via the *GeneSwitch* system. In the order of **i** to **p**, *Control* flies used are 134, 87, 125, 102, 106, 122, 100 and 120; flies used in the experimental groups were 129, 80, 87, 122, 114, 77, 48 and 129. **q**,**r**, quantifications of gut mitoses for flies with the indicated genotypes raised on CD for 20 days. Each dot stands for one gut. Data are mean ± s.e.m. in **b**,**d**,**e**,**q**,**r** with Student’s *t*-test. Log-rank test in **f**-**p**. *P* values are indicated.

**Extended Data Fig. 3.**
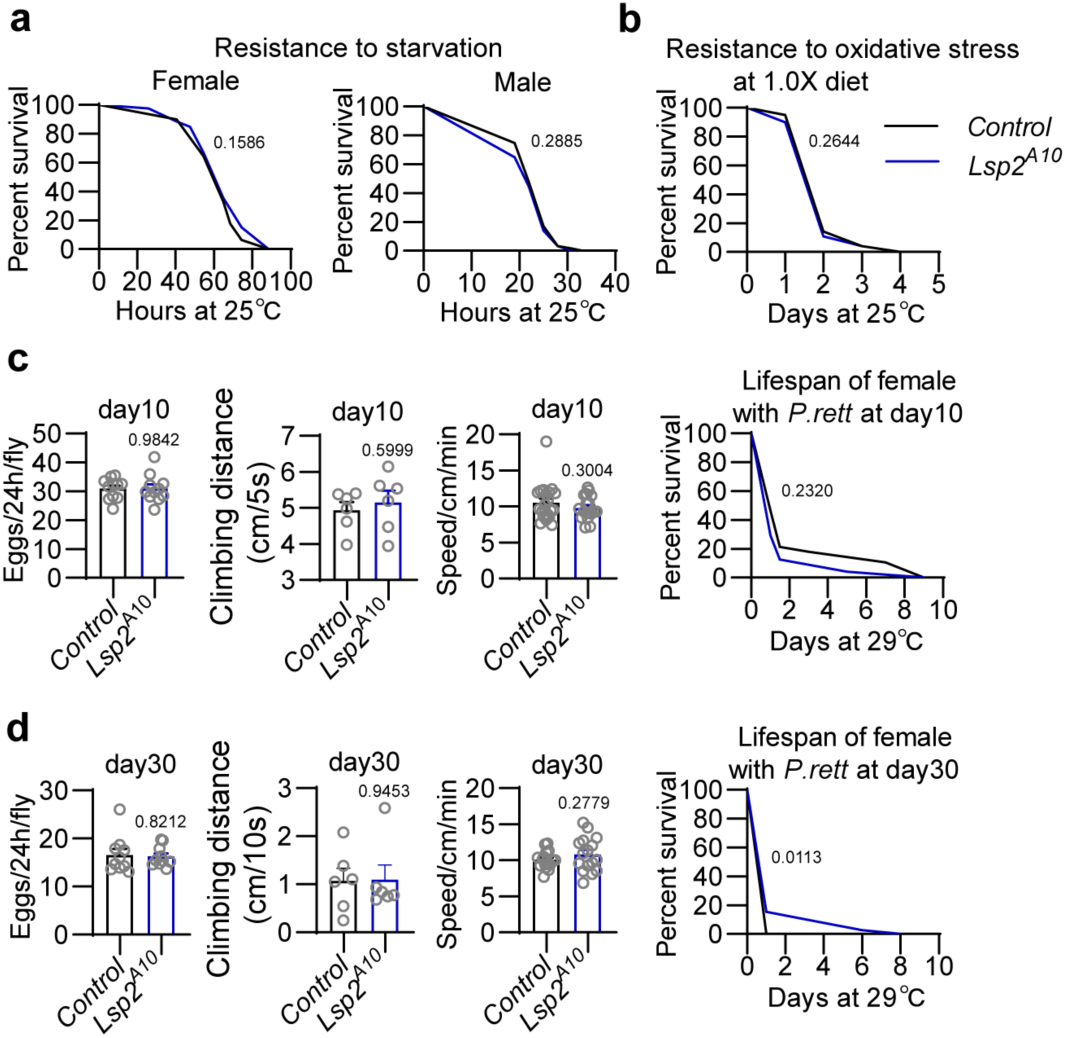
Loss of *Lsp2* does not alter stress resistance, locomotor activity and reproduction. **a**, Survival of *Control* and *Lsp2^A10^* flies to starvation. 5-day-old flies were analysed on 1% agar tube with only the access to water. For female (*left*), *n =* 79 in both *Control* and *Lsp2^A10^*; for male (*right*), *n =* 122 in *Control* and n = 128 in *Lsp2^A10^*. **b**, Survival of 5-day-old *Control* and *Lsp2^A10^* females to oxidative stress. 17.5μM paraquat was applied to 1.0X SYA food to feed flies. *n =* 120 for both *Control* and *Lsp2^A10^*. **c**,**d**, Eggs laid per fly per 24 hours (*left*), climbing distance in the indicated time (*left in the middle*), average speed of spontaneous movements monitored for 8 hours (*right in the middle*) and survival to systemic bacterial infection of *Providencia rettgeri* (*P. rett*) (*right*) in 10-day-old (**c**) and 30-day-old (**d**) *Control* and *Lsp2^A10^* females. For egg laying, one dot represents a group of 5 females and 3 males; for climbing, one dot stands for the average distance climbed by a group of 10 females; for spontaneous walking, one dot represents one fly; for infection experiments, flies used for *Control* and *Lsp2^A10^* are 28 and 24 in **c**, and 39 and 39 in **d**. Data are mean ± s.e.m. except survival plots. Student’s *t*-test for bar plots; Log-rank test for survival analyses. *P* values are indicated.

**Extended Data Fig. 4.**
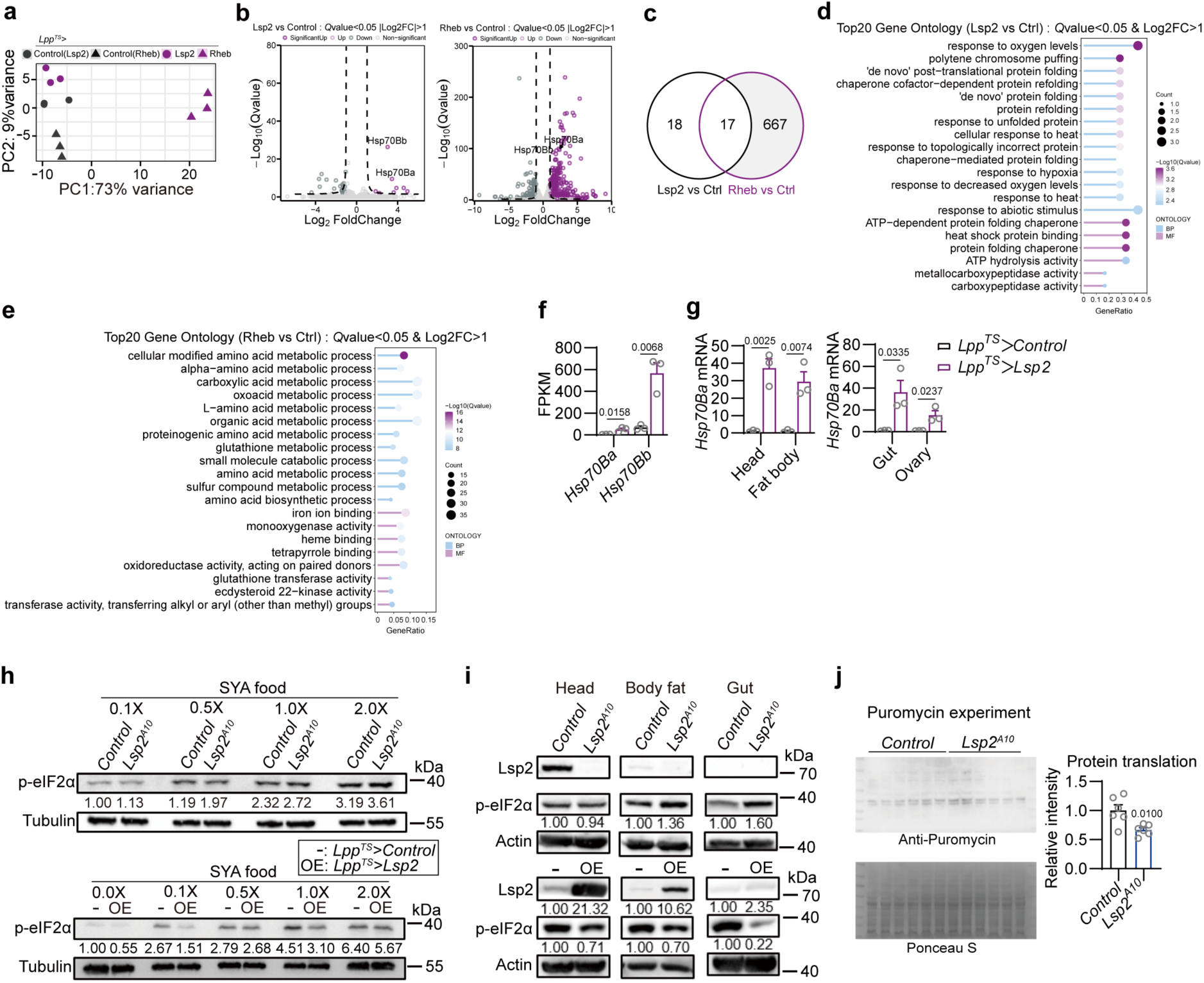
Implications for Lsp2 in regulating translation. **a-e**, RNA-seq analyses suggest Lsp2 is not a transcriptional regulator. PCA analysis (**a**) of differentially expressed genes (DEGs, *Q*<0.05 & |log_2_FoldChange|>1; volcano plots shown in **b**) in heads of adult females with the indicated genotypes raised at 29℃ for 5 days (3 days on conventional lab diet (CD) followed by 2 days on 0.0X SYA food) to overexpress *Lsp2* or *Rheb* in the fat body using *Lpp^TS^*. Shown in **c** are shared DEGs between the two datasets, including *Hsp70Ba* and *Hsp70Bb* also highlighted in **b**. Gene ontology (GO) analyses of upregulated DEGs found in each dataset (**d**,**e**). **f**,**g**, *Hsp70Ba* and *Hsp70Bb* expression levels in the RNA-seq data (**f**) and qPCR (**g**) detecting the relative *Hsp70Ba* mRNA levels in the indicated tissues (head, fat tissue in the body, gut and ovary) of female flies of *Lpp^TS^>Control* and *Lpp^TS^>Lsp2* raised at 29℃ for 5 days (3 days on CD followed by 2 days on 0.0X SYA food). *Hsp70Ba* expression in tissues of *Lpp^TS^>Control* fly is normalized to 1. One dot stands for a group of 20 flies. **h**, Western blots measuring levels of phosphorylation of the eukaryotic initiation factor 2 (eIF2) α subunit (p-eIF2a) in 5-day-old flies with the indicated genotypes raised on SYA diets for 7 days. Whole flies were used for blotting. **i**, Western blots measuring levels of Lsp2 and p-eIF2a in heads, fat tissue in the body and guts of 5-day-old flies with the indicated genotypes raised on CD for 7 days. *Lpp^TS^>Lsp2* is abbreviated as OE in **h**,**i**. **j**, Protein synthesis in *Control* and *Lsp2^A10^* flies and its quantification. 3-day-old females were raised on CD for 7 days followed by a 1.0X SYA diet containing 600 μM puromycin for 24 hours. Whole flies were analysed. Ponceau S staining indicates total protein. Each dot stands for a group of 20 flies. Student’s *t*-test in **f**,**g**,**j** with *P* values indicated.

**Extended Data Fig. 5.**
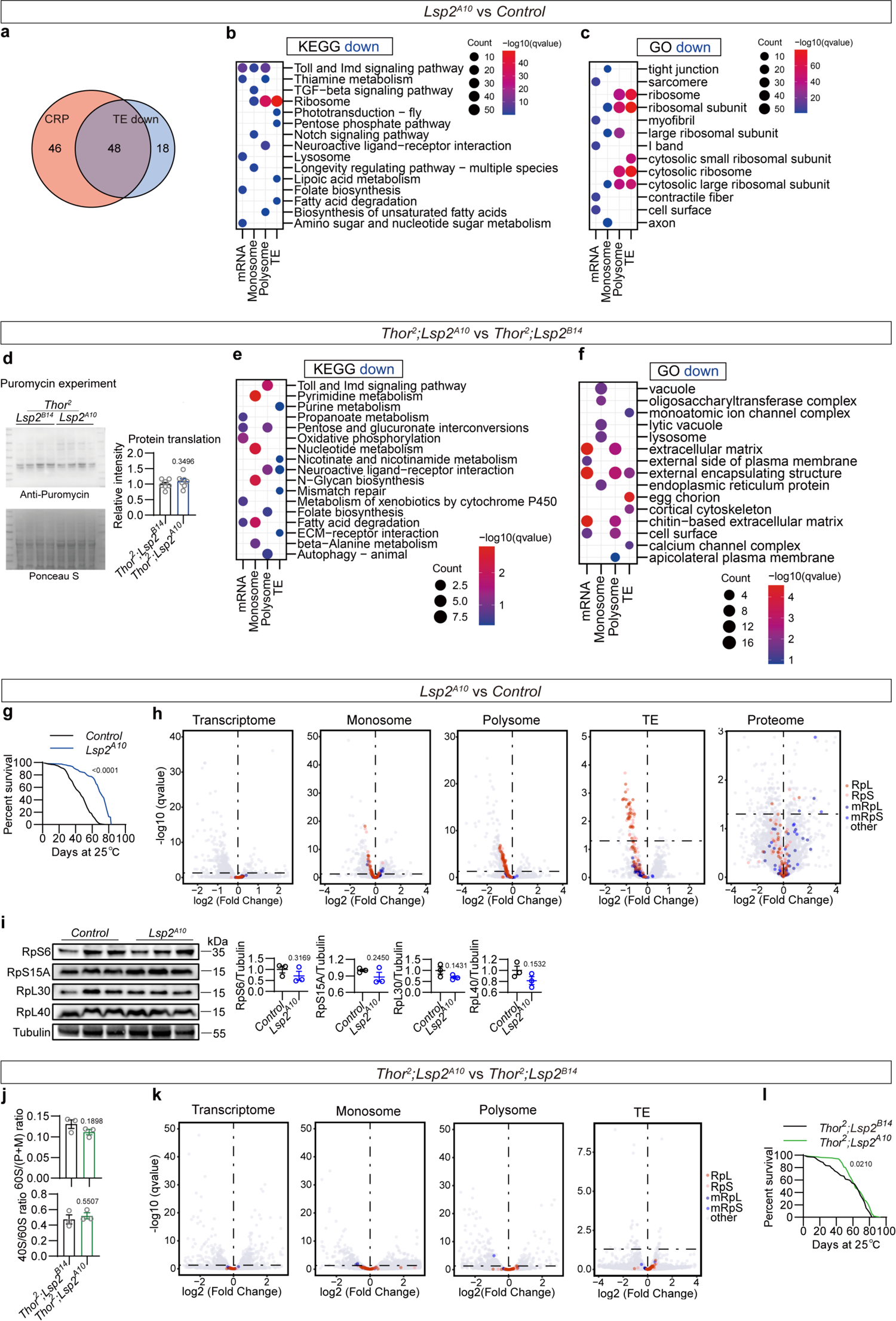
Lsp2 impacts lifespan by regulating the translation of RP mRNAs through 4E-BP. **a**, Overlap between fly RP genes (94 in total) and translationally downregulated genes (66) by *Lsp2* loss. **b**,**c**, KEGG (**b**) and Gene ontology (GO; **c**) analyses of downregulated DEGs in *Lsp2^A10^*flies compared to *Control* in levels of mRNA-seq, monosome-seq, polysome-seq and translational efficiency (TE). Shown are the top 5 enriched processes in each level. **d**, Protein synthesis in *Thor^2^; Lsp2^B14^* and *Thor^2^; Lsp2^A10^* flies and its quantification. 3-day-old females were raised on conventional lab diet (CD) for 7 days followed by a 1.0X SYA diet containing 600 μM puromycin for 24 hours. Whole flies were analysed. Ponceau S staining indicates total protein. **e**,**f**, KEGG **(e**) and GO (**f**) analyses of downregulated DEGs in *Thor^2^; Lsp2^A10^* flies compared to *Thor^2^; Lsp2^B14^*flies in levels of mRNA-seq, monosome-seq, polysome-seq and TE. Shown are the top 5 enriched processes in each level. **g**, Lifespan of *Control* (n = 127) and *Lsp2^A10^* (n = 124) females raised on CD. **h**, Volcano plots of expression changes in mRNA-seq, monosome-seq, polysome-seq, TE and proteomics in *Lsp2^A10^* mutant flies compared to *Control* flies. RP and mitochondrial RP (mRP) mRNAs/proteins are highlighted. The dashed horizontal lines denote thresholds. **i**, Western blots and the quantification of the indicated RP proteins in whole flies of *Control* and *Lsp2^A10^* lines raised on CD for 7 days. **j**, Polysome profiling of 15-day-old *Thor^2^; Lsp2^B14^* and *Thor^2^; Lsp2^A10^* flies raised on 0.5X SYA food reveals similar levels of the 60S subunit (*upper*) and the 40S subunit (*bottom*). **k**, Volcano plots of expression changes in mRNA-seq, monosome-seq, polysome-seq and TE in *Thor^2^; Lsp2^A10^* double mutant flies compared to *Thor^2^; Lsp2^B14^* flies. RP and mRP mRNAs are highlighted. The dashed horizontal lines denote thresholds. **l**, Lifespan of *Thor^2^; Lsp2^B14^* (n = 127) and *Thor^2^; Lsp2^A10^* (n = 116) females raised on CD. Each dot stands for a group of 20 flies in **d**,**i**. Data are mean ± s.e.m. in **d**,**i**,**j** with Student’s *t*-test. Log-rank test in **g**,**l**. *P* values are indicated.

**Extended Data Fig. 6.**
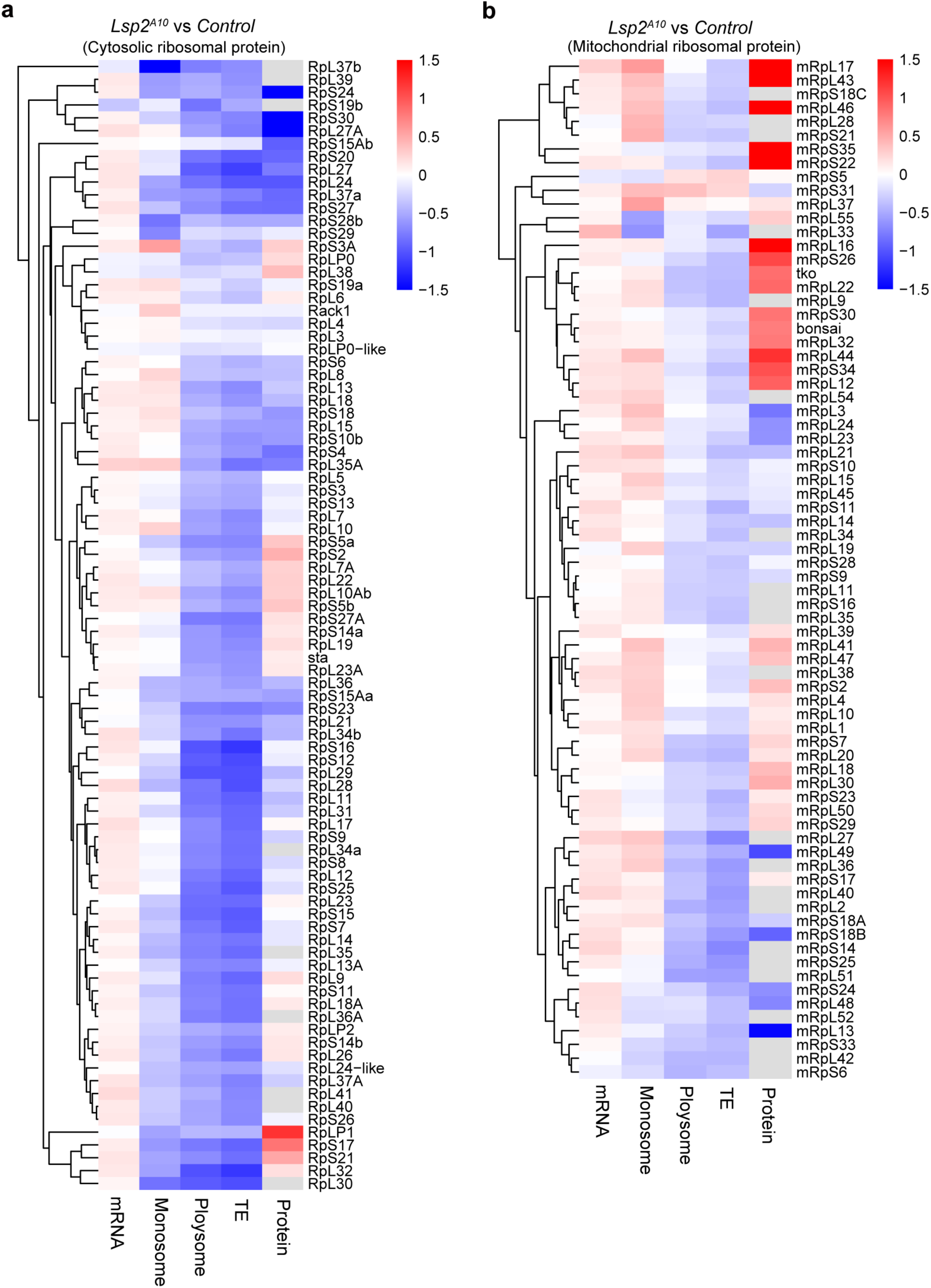
Translational changes in RP and mRP genes in *Lsp2* mutants. **a**,**b**, Heatmaps showing expression changes in RP (**a**) and mRP (**b**) mRNA/protein levels in mRNA-seq, monosome-seq, polysome-seq, TE and proteomics in *Lsp2^A10^* mutant flies compared to *Control* flies.

**Extended Data Fig. 7.**
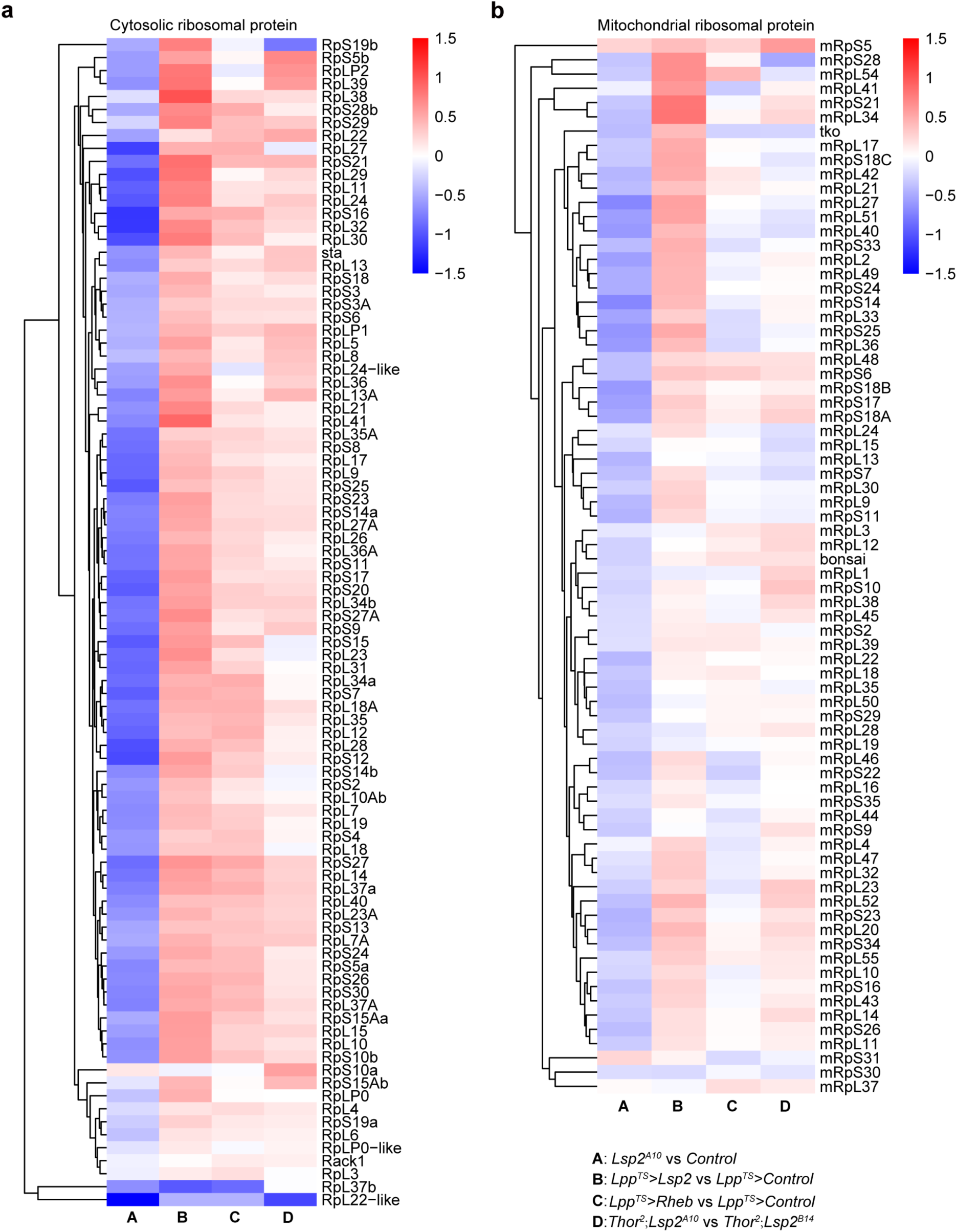
Lsp2 regulates the translational efficiencies of RP but not mRP mRNAs in a 4E-BP-dependent manner. **a**,**b**, Heatmaps summarising changes in translational efficiencies of RP (**a**) and mRP (**b**) mRNAs in the four indicated datasets generated in this study.

**Extended Data Fig. 8.**
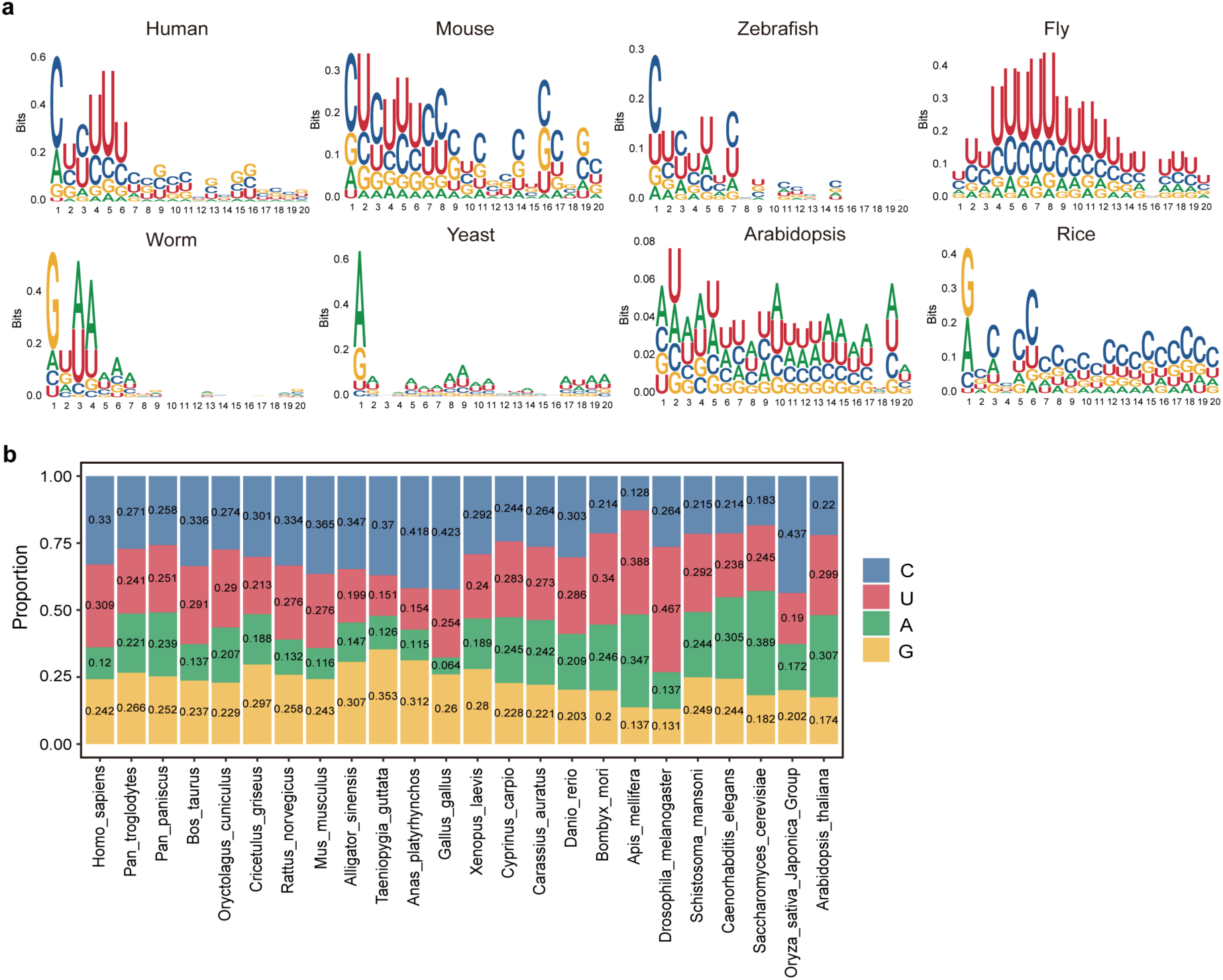
Identification of TOP motifs in RP mRNAs across representative species. **a**, Sequence logos of the first 20nt of 5’ UTR of 8 representative species. **b**, Stacked bar chart of 5’ UTR sequences from RP mRNAs of 24 species.

**Extended Data Fig. 9.**
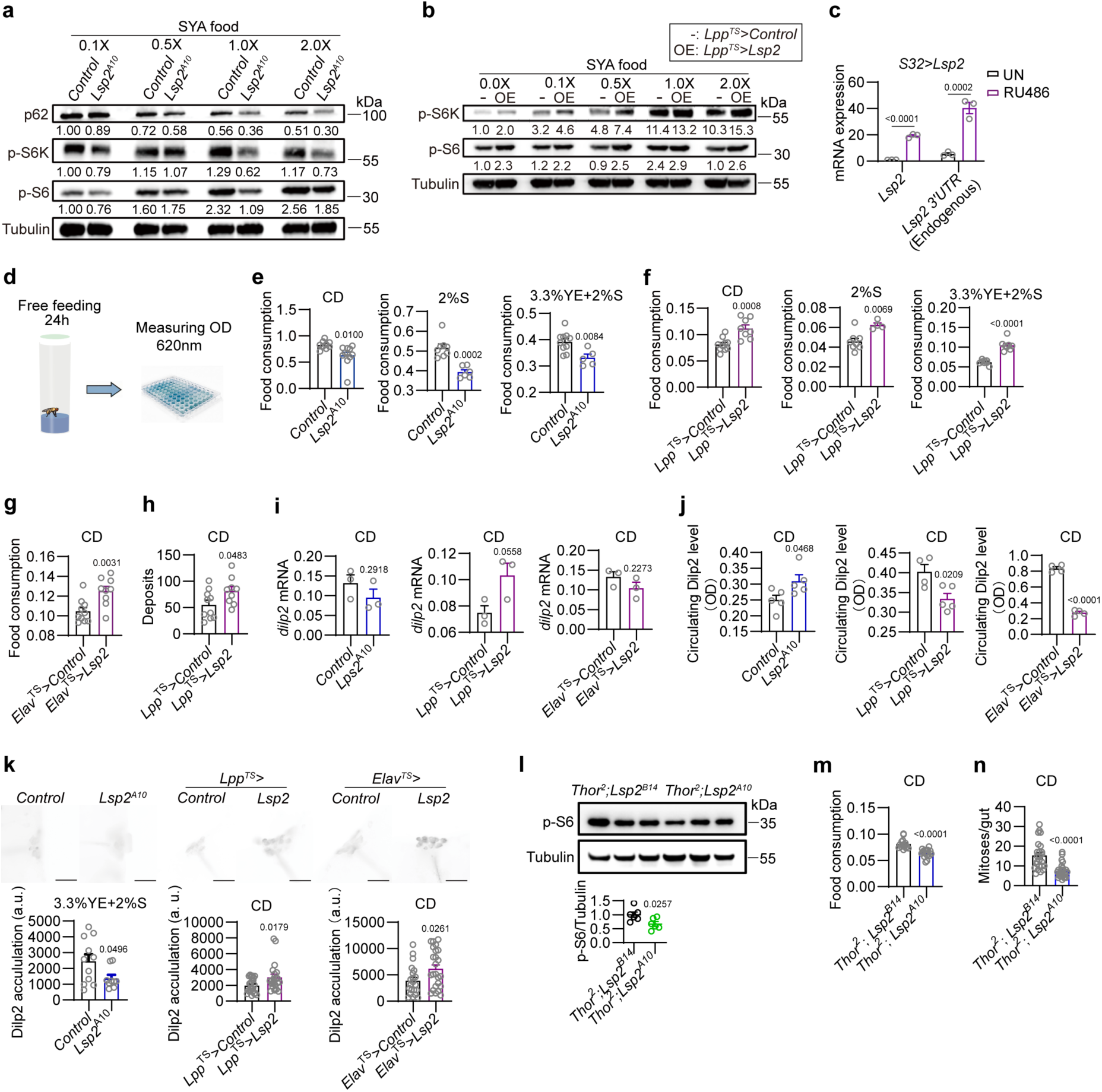
Lsp2 promotes mTORC1 activity and food intake independently of 4E-BP. **a**,**b**, Western blots quantifying the phosphorylation of S6K on Thr398 (p-S6K, a canonical mTORC1 readout), its product phospho-S6 ribosomal protein (p-S6), and p62 (Ref(2)P in *Drosophila*) in 5-day-old flies with the indicated genotypes raised on SYA diets for 7 days. Whole flies were used. Note that p62 marks ubiquitinated protein bodies for autophagic degradation and accumulates in cells during inhibition of autophagy. The results show that *Lsp2* loss (**a**) reduces while *Lsp2* overexpression (**b**) increases these canonical mTORC1 activity indicators. **c**, Overexpressing *Lsp2* increases the transcription of endogenous *Lsp2* gene (measured with primers targeting 3’ UTR of *Lsp2*), supporting that Lsp2 promotes mTORC1 activity thereby forming a feedback amplification loop to sustain its robust upregulation by EAAs. Shown are qPCR assays detecting regions of *Lsp2* CDS and 3’ UTR in heads of *S32-GS-Gal4>Lsp2* flies with/without adding RU486 to induce transgene expression for 3 days on conventional lab diet (CD). **d**, A schematic of food intake measurement using colorimetric assay. **e**-**g** Food consumption of 5-day-old females with the indicated genotypes on CD, 2% sucrose and a diet consisting of 3.3% YE+2% sucrose. *Elav-Gal4* is a pan-neuronal driver. One dot denotes a group of 15 females. The changes in food intake were consistently observed in CD, sugar-only diets, and protein diets, suggesting that Lsp2 does not drive nutrient-specific appetite. **h**, Food consumption by measuring excreta of 5-day-old *Lpp^TS^>Control* and *Lpp^TS^>Lsp2* females. One dot denotes deposits over 24 hours by a group of 5 females. **i**-**k**, qPCR measuring *Drosophila* insulin-like peptide 2 (*Dilp2*) mRNA levels (**i**), ELISA quantification of circulating Dilp2 protein levels (**j**) and immunofluorescence showing levels of Dilp2 retaining in the insulin-producing cells (IPCs) (**k**) of flies with the indicated genotypes raised on CD except the *Control* and *Lsp2^A10^* in **k**. Heads (**i**), haemolymph (**j**) and brains (**k**) of 5-day-old females were used. One dot in **i** stands for a group of 20 flies, in **j** for 50 flies and one dot in **k** denotes one brain. The results support that Lsp2 adipokine acts to limit the release of brain insulin Dilp2 thereby promoting food intake. **l-n**, Like *Lsp2^B14^*and *Lsp2^A10^* flies, *Thor^2^; Lsp2^B14^*and *Thor^2^; Lsp2^A10^* flies still differ in p-S6 levels (**l**), food intake (**m**) and division of intestinal stem cells (**n**). 7-day-old whole flies raised on CD were used in **l**, and one dot denotes a group of 20 females. For food intake in **m**, one dot denotes a group of fifteen 7-day-old females raised on CD. For stem cell division in **n**, one dot stands for one gut of a 10-day-old female raised on CD. Data are mean ± s.e.m. for all the bar plot quantifications. Student’s *t*-test. *P* values are indicated. Scale bars 50 μm in **k.**

**Extended Data Fig. 10.**
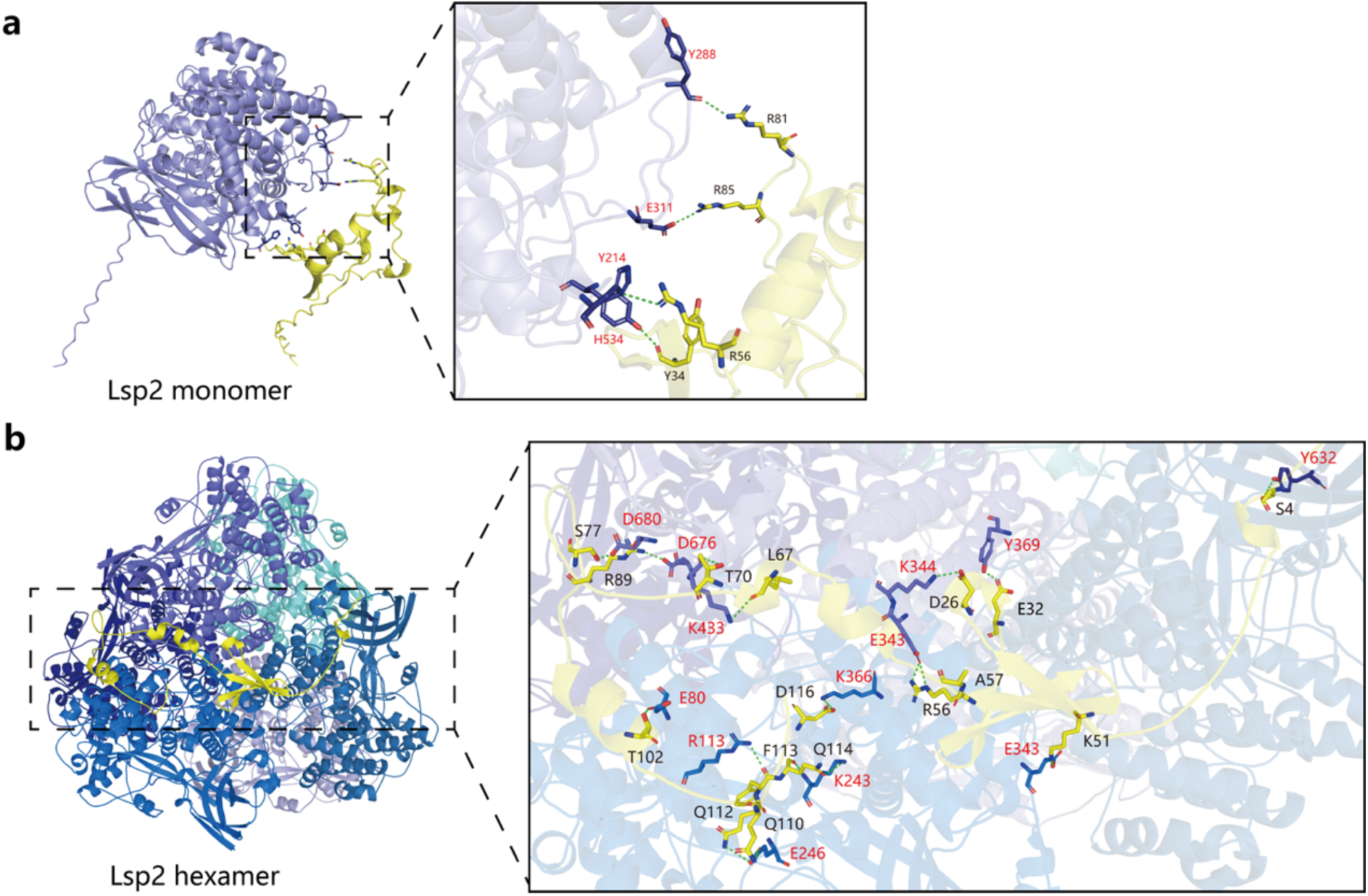
Structural modeling predicts interactions between Lsp2 and 4E-BP (Thor). **a**, Predicted structural model of the interaction between Thor and the Lsp2 monomer. *Left*: The Lsp2 monomer is shown in purple, and the Thor protein is depicted in yellow using both cartoon and ribbon representations. *Right*: Close-up view of the predicted interaction interface between the Lsp2 monomer (purple) and Thor (yellow). **b**, Predicted structural model of the interaction between Thor and the Lsp2 hexamer. *Left*: The Lsp2 hexamer is shown in purple, cyan, and blue, while Thor is displayed in yellow. *Right*: Close-up of the interaction interface, highlighting 17 predicted hydrogen bonds (green dashed lines), including the phosphorylatable residue Thr70 of Thor, a phosphorylation target of mTORC1. Residue labels for Lsp2 are shown in red, and those for Thor in black.

**Extended Data Table 1.**
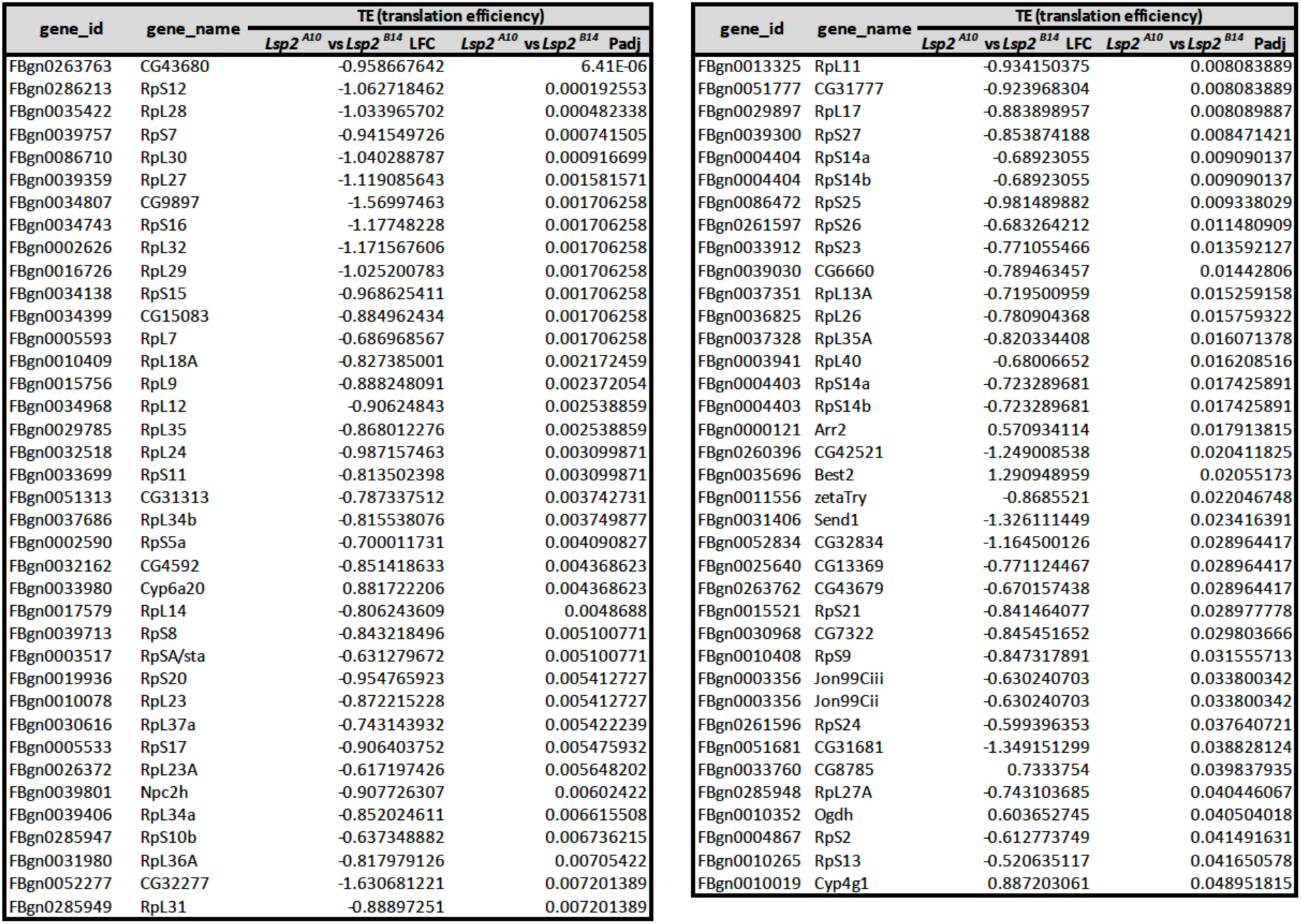
Genes with significantly altered TEs in *Lsp2^A10^* flies.

**Extended Data Table 2.**
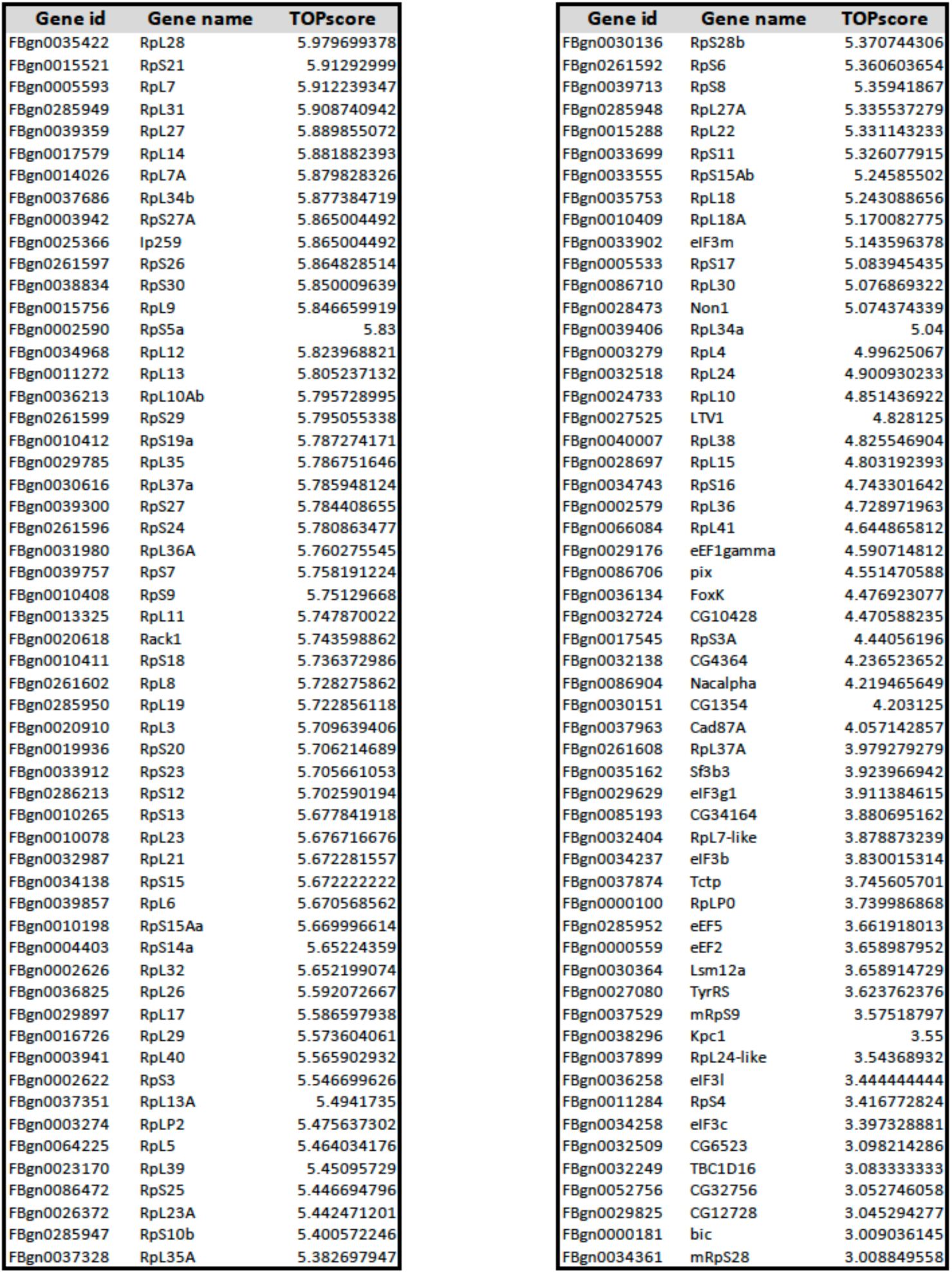
Predicted *Drosophila* TOP genes and their TOP scores.

**Extended Data Table 3.**
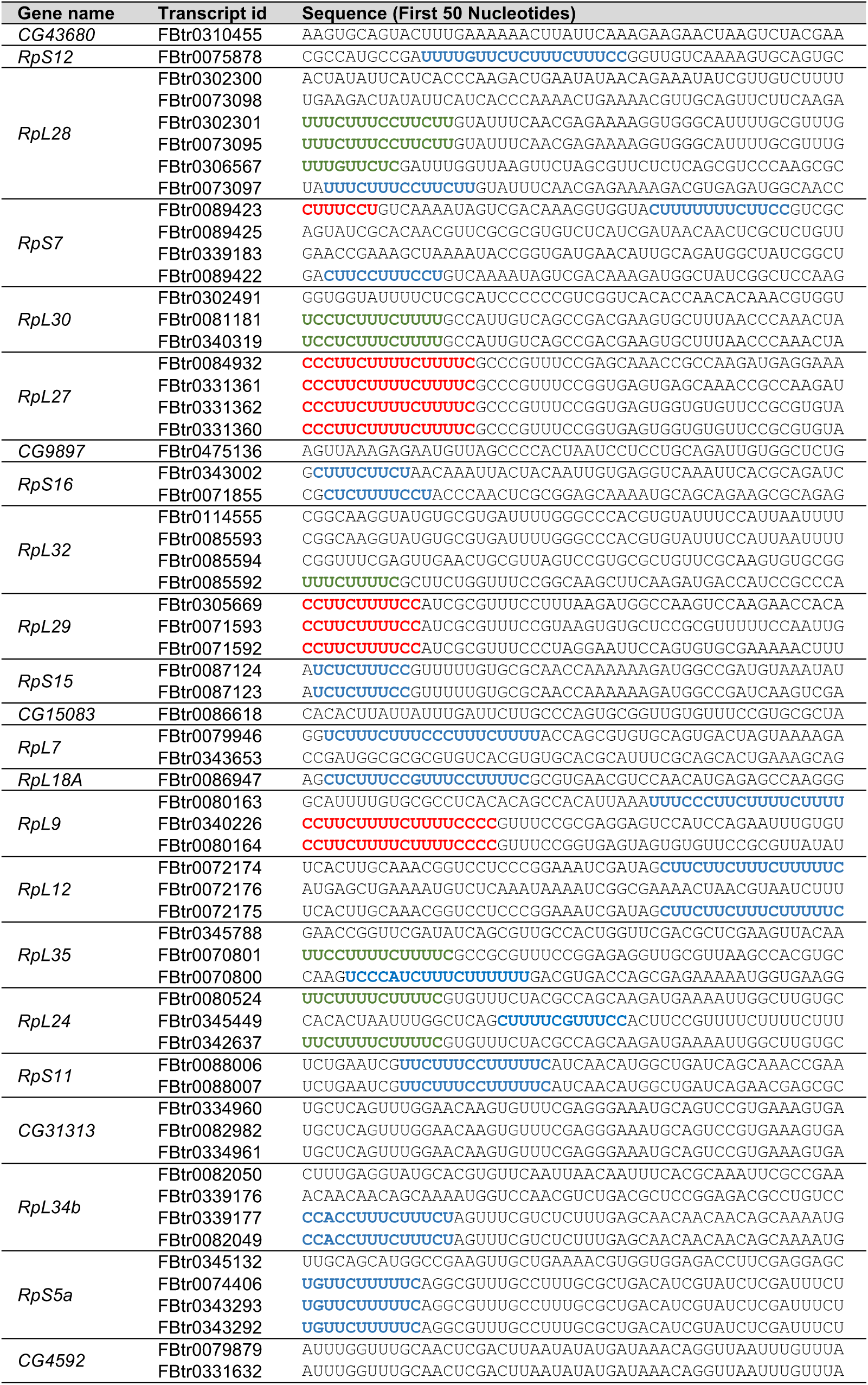
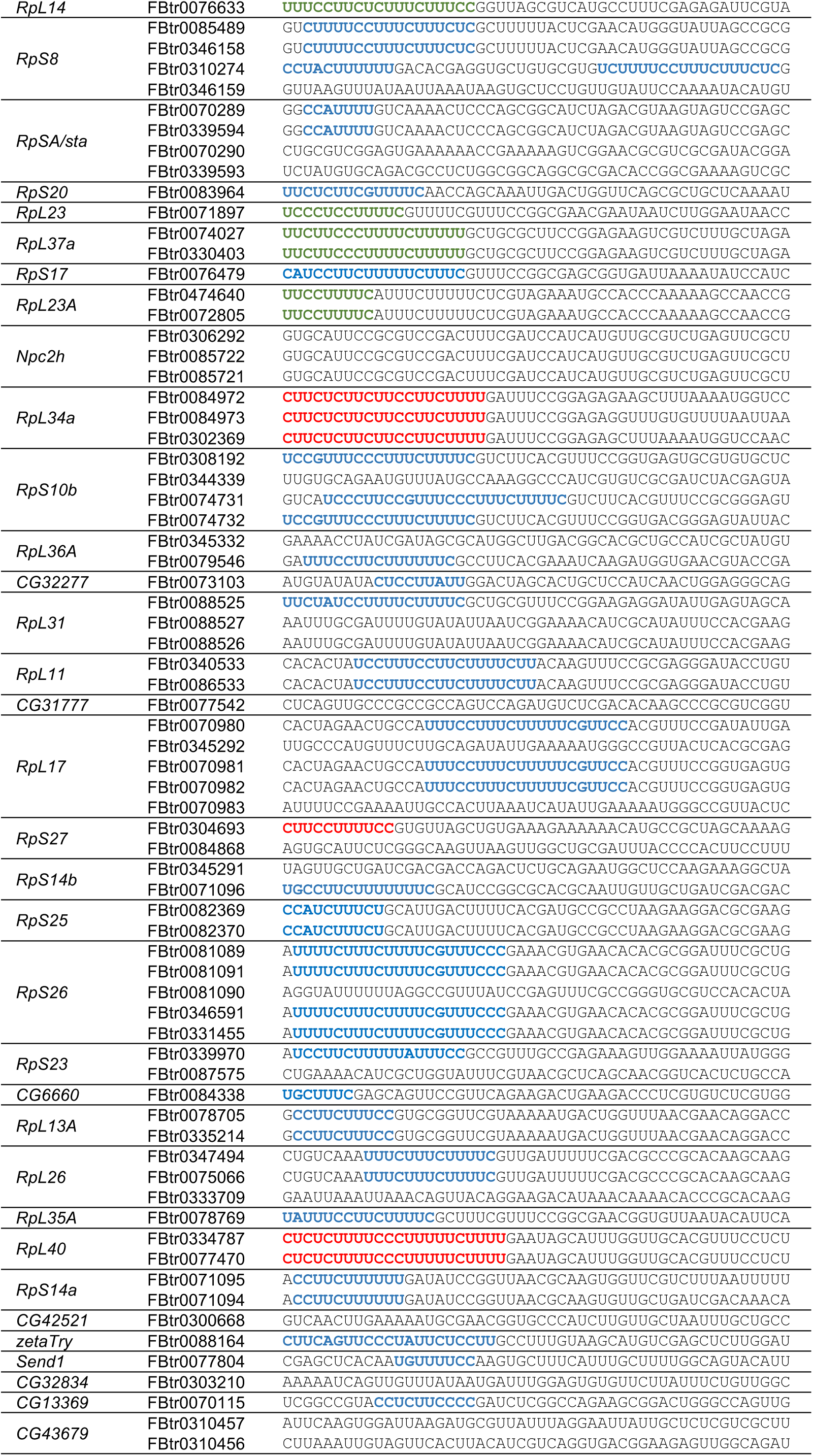
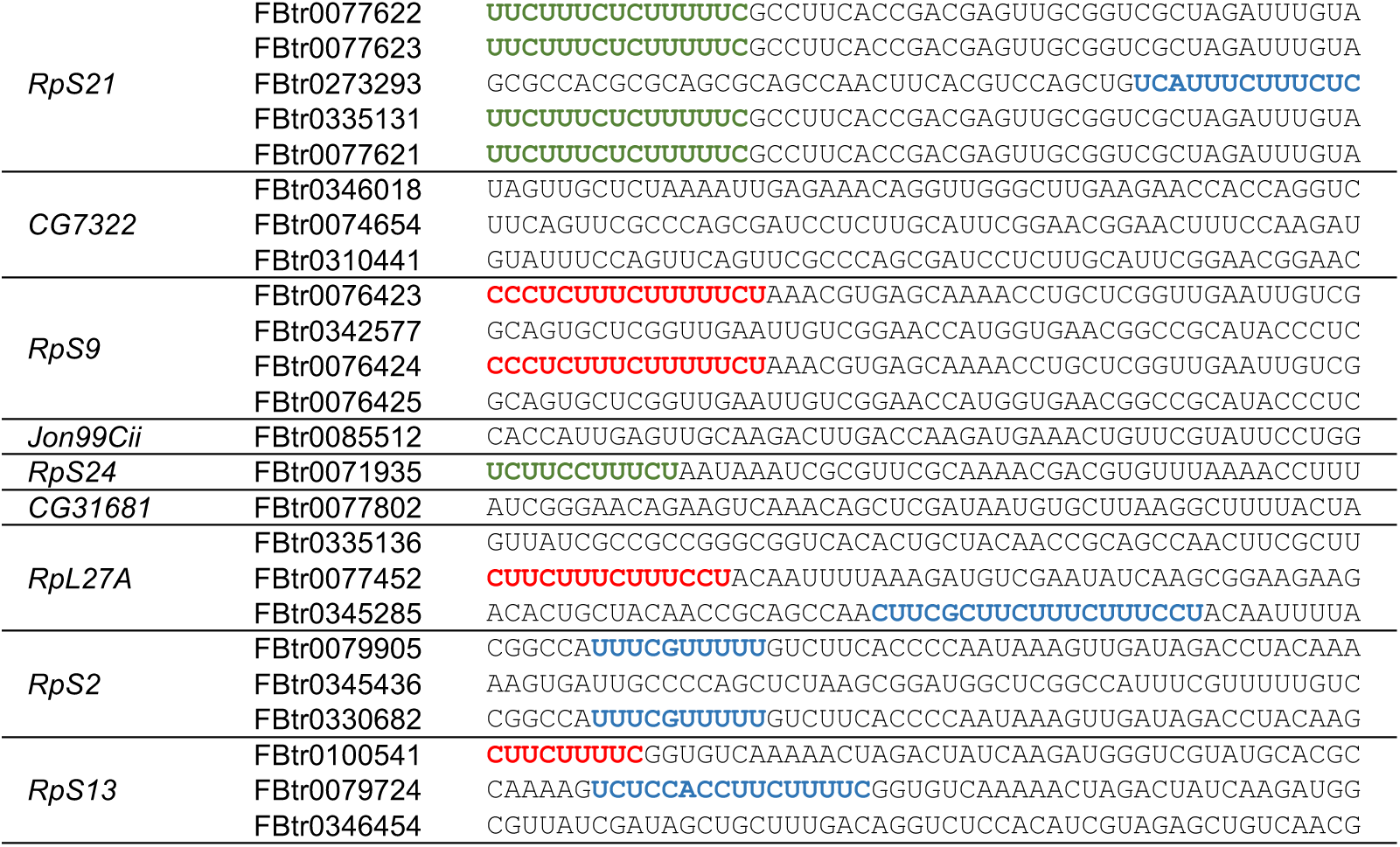
Fly TOP motifs found in TE-downregulated genes in *Lsp2^A10^* flies. Canonical TOP motifs are marked in red, iso-TOP motifs in green and pyrimidine-rich sequences in blue. These three types of colored motifs, collectively termed as fly TOP motifs, are candidates that confer translational control by the mTORC1 nutrient-sensing pathway.

